# Pronounced Sex Differences in Evoked and Spontaneous Pain Assessments Following Full-Thickness Traumatic Burn Injury in Male and Female Sprague Dawley Rats

**DOI:** 10.64898/2026.03.12.711381

**Authors:** Corinne M. Augusto, Ann Sipe, Cole F. P. Moran-Bariso, Charles N. Zawatsky, Jennifer E. Nyland

## Abstract

Persistent pain is a common but poorly understood outcome of traumatic burn injury. With increasing numbers of patients surviving their burn injuries, ongoing pain presents a growing complication to patient healing and quality of life. Despite more women reporting chronic pain post-burn than men, preclinical burn research rarely includes female animals. To address this gap, this study examined a diverse set of behavioral outcomes in male and female rats after a unilateral full-thickness burn to the hind paw. Utilizing traditional methods to assess evoked pain behaviors, new technology to assess gait abnormalities, and established techniques to evaluate comorbid anxiety-like behavior, we determined that male and female rats have divergent pain-related behaviors post-burn. Both sexes experienced mechanical allodynia after burn injury, but only males experienced thermal hyperalgesia. In contrast, female rats were acutely resistant to noxious thermal stimulation. While both sexes demonstrated gait abnormalities post-burn when freely ambulating, female rats exhibited a wider range of abnormal gait features, which were more severe and longer-lasting than those in males. However, despite both sexes demonstrating symptoms of persistent pain, only males displayed anxiety-like behavior on the Elevated Zero Maze. In conclusion, our study found that male and female Sprague Dawley rats displayed divergent, sex-specific evoked pain responses, gait dysfunction, and anxiety-like behavior after full-thickness burn injury. Future studies should examine the underlying mechanisms behind these behavioral sex differences.

**Perspective:** This article takes a novel approach to pain behavior testing after full-thickness burn injury, capturing behaviors beyond traditional reflexive (“evoked”) behaviors. The results of this article provide evidence that preclinical research must expand behavioral testing to capture the full animal pain experience and better model human patient outcomes.

## Introduction

The WHO estimates 180,000 people worldwide die annually from burns,^1^ with hundreds of thousands of burn incidents occurring annually in the U.S. alone; there were an estimated 404,000 emergency department visits for burns or corrosion in 2021,^2^ and 5,505 deaths occurred from fire, heat, or hot substances in 2017.^3^ The American Burn Association (ABA) reports that 97.6% of patients admitted to burn centers between 2020 and 2024 survived.^4^ In light of the tremendous survival rate, non-fatal burns are a leading cause of morbidity, disfigurement, and disability, and typically require prolonged hospitalization.^1^ Acute burn pain can be excruciating, particularly during dressing changes or debridement procedures.^5,6^ Patients with burns greater than 25% total body surface area (TBSA) are at greater risk of systemic injury, including but not limited to: pulmonary edema, hypercoagulability, bronchoconstriction, and hypermetabolism.^7^ Additionally, burn injuries leave patients vulnerable to chronic pain and comorbid psychopathology.

The incidence of chronic pain months to years post-burn varies widely – anywhere from 6% to 36% of patients report lasting pain.^8–13^ Patients with severe (> 20% TBSA) burns are more likely to report moderate or severe pain (average of 5 months post-discharge) compared to matched normative controls.^14^ Burn severity, depressive symptoms, age, substance use, prolonged hospitalizations, hypertrophic scar severity, and severe medical interventions (i.e., intubation, surgeries) are common predictors of post-burn pain severity.^8,10,11^ The likelihood of chronic pain post-burn is also affected by the anatomical location of the burn(s), with upper arm burns and thigh burns being the strongest anatomical predictors of chronic pain.^15^ Importantly, chronic pain post-burn is associated with worse physical and emotional functioning.^9^

Patients with burns > 20% TBSA have been shown to be more likely to report pain interference of any severity, and less likely to report the absence of anxiety or depressive symptoms compared with normative controls.^14^ Patients often experience “breakthrough” pain, a transient increase in their pain intensity, often associated with movement (but can also be spontaneous).^16^ Unfortunately, many of these spontaneous pain experiences are not assessed in preclinical research. To improve the translational relevance of our results, this study employed diverse techniques to assess pain, moving beyond classical evoked-pain methods that capture symptoms like mechanical allodynia and thermal hyperalgesia. While the evoked-pain tests are the gold standard for detecting persistent pain in rodent models, this study used additional techniques to assess comorbid psychological distress (e.g., the Elevated Zero Maze [EZM] for anxiety) and reduced physical functioning (e.g., detecting gait abnormalities in voluntary ambulation).

Burn injuries expose patients to psychopathology both during and after burn injury, creating a negative feedback loop with pain outcomes. In a study using TriNetX data, patients with severe (40-59% TBSA) or critical (> 60% TBSA) burns had an incidence of post-traumatic stress disorder (PTSD) of 12.26% and 10.81%, respectively, at least 1-month post-burn.^17^ Of the data examined, patients with comorbid PTSD had increased risk of anxiety, opioid use disorder (OUD), chronic pain, and/or depression compared to TBSA-matched patients without PTSD.^17^ PTSD severity at 6 months has also been shown to correlate positively with current graft site itch and pain.^11^ To increase the translational relevance of our results, this study used a rodent burn model involving a unilateral full-thickness burn injury to the plantar surface of one hind paw, preceded by peri-injury stressors to replicate both the physical *and* psychological trauma typically experienced by humans, as described previously.^18^

Women are more likely to report higher levels of pain and are more likely than men to suffer from chronic pain conditions.^19,20^ In one study of major thermal burn injuries, 32% of women compared with 5% of men had severe chronic pain one year after injury.^21^ The study also found that women experienced greater pain severity at 3, 6, and 12 months after the injury. Despite female sex being identified as a risk factor for pain, depression, and anxiety post-burn in clinical studies,^14^ female rats and mice have not historically been used in preclinical research. Over the last decade, increased inclusion of female subjects has suggested that male and female rats differ in their mechanical sensitivity to burn injury;^22,23^ however, direct comparisons have not been conducted, nor have assessments of non-evoked pain. As such, our study utilized equal numbers of gonadally-intact male and female Sprague-Dawley rats and assessed both evoked and spontaneous pain behaviors.

## Methods

### Animals & Group Assignments

This study used 24 male and 24 female Sprague-Dawley rats (Charles River, Wilmington, MA), which arrived at approximately 70 days old. All rats were single-housed in solid-bottom cages with TEK-fresh bedding (Envigo Bioproducts, Inc., Indianapolis, IN), and *ad libitum* access to food and water. Rats were housed in a temperature- and humidity-controlled facility with a 12-hour light/dark cycle, 72°F ambient temperature, and 40% relative humidity. Rats were allowed one week to acclimate to the facilities and another week to be handled by the investigators before experimental procedures began. The housing environment and cohort health were monitored daily by trained animal care technicians employed by the Department of Comparative Medicine. All experimental procedures were approved by the Penn State College of Medicine Institutional Animal Care and Use Committee (PROTO202202209) and were conducted in accordance with the National Institutes of Health (NIH) Guide for the Care and Use of Laboratory Animals. All efforts were made to minimize animal suffering and the number of animals used.

### Experimental Timeline

Upon arrival, rats acclimated to the facility for 1 week (days -18 to -12), then were gentled by lab personnel for 5 days (days -11 to -7). After this gentling period, all rats underwent baseline assessments of spontaneous pain behaviors, evoked pain responses, and alternative pain behavior testing (i.e., EZM). As previously described,^18^ the traumatic burn injury model consists of three sessions of forced swim stress followed by induction of a thermal injury to one hind paw. Following baseline testing, rats were randomly assigned to the experimental or control condition and underwent three consecutive days of forced swim stress or sham swim stress. Spontaneous, evoked, and alternative pain behaviors were retested 24 hours following the final swim stress or sham stress exposure, followed by induction of a full-thickness thermal injury to the right hind paw, or sham injury for those in the control condition. This resulted in 12 male and 12 female rats experiencing both the swim stress and the burn injury (“Burn” rats), while 12 male and 12 female rats experienced the sham swim stress and sham injury (“Control” rats). Evoked pain (EP) and spontaneous pain (SP) behaviors were retested 1, 3, 7, 14, 21, 28, 35, 42, 49, and 56 days post-injury. EZM was reassessed at 10, 24, 38, and 52 days post-injury. All behavioral testing was performed during the light cycle. After the final behavioral test, all rats were humanely euthanized. See **Figure 1** for a visual timeline.

**Figure 1.**
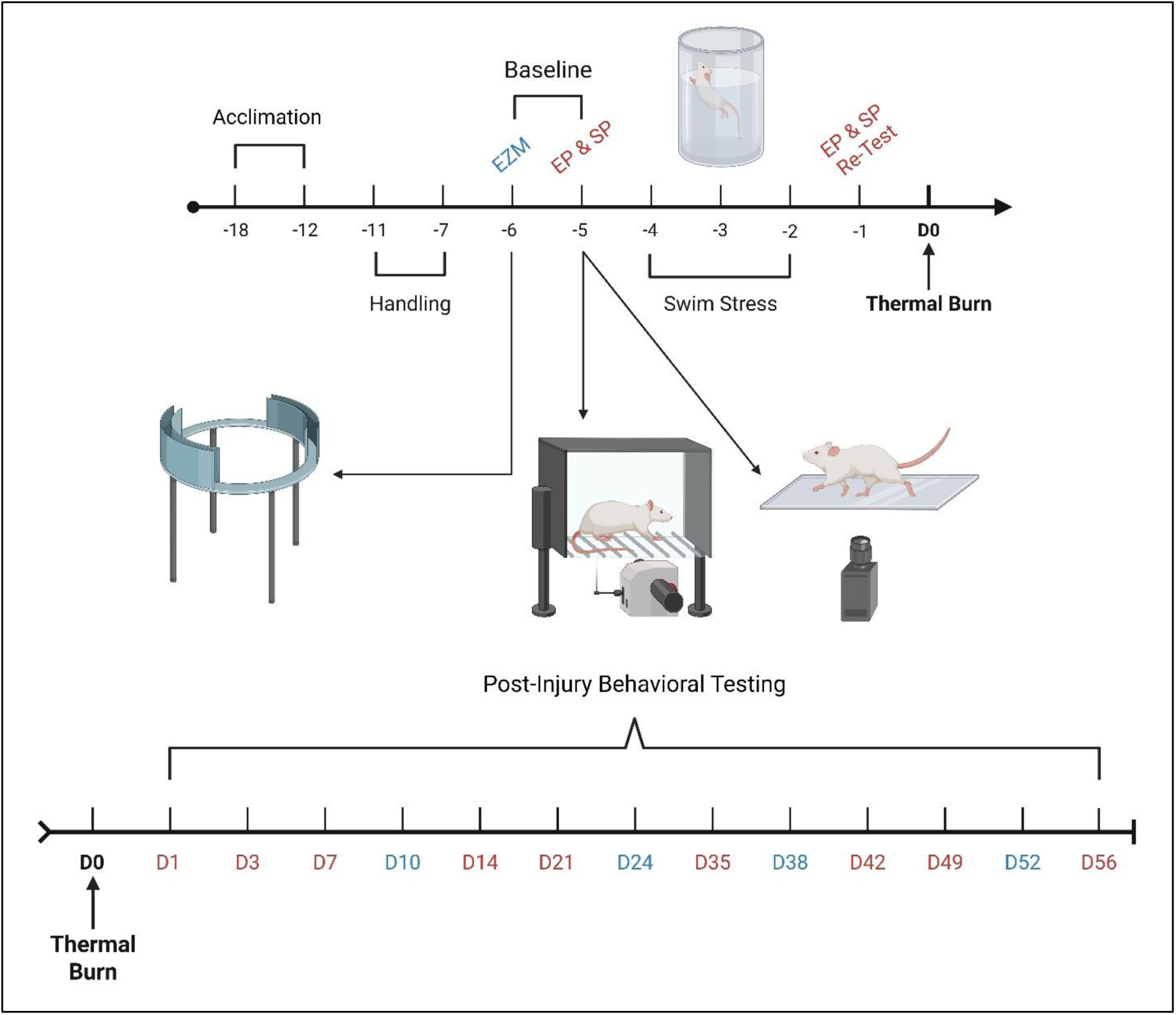
Experimental Timeline. Red font denotes Evoked and Spontaneous Pain behavior testing. Blue font denotes Elevated Zero Maze testing. *EZM*, Elevated Zero Maze; *EP*, Evoked Pain behavior testing; *SP*, Spontaneous Pain behavior testing. Figure created in BioRender

### Swim Stress Procedure

This procedure was derived from previously published work.^18^ Briefly, rats were individually placed in translucent Plexiglass cylinders (30 cm diameter x 50 cm height; Panlab). For the swim stress condition, these cylinders were filled with room temperature water (17.8-19.5 °C) to a depth of 20 cm. This depth has been confirmed to be deep enough to force rats to swim as they cannot reach the bottom of the cylinder with their toes or tails.^18^ For the sham stress condition, the cylinders were left dry and empty (i.e., no bedding). All rats underwent daily 20-minute swim or sham stress sessions for three consecutive days. After each swim stress session, experimenters dried the wet rats with clean microfiber towels, rubbing vigorously until the rats’ fur was no longer waterlogged; rats were returned to home cages once dry. All cylinders (wet & dry) were cleaned of excrement between rats, and with 70% ethanol between testing days.

### Thermal Injury & Sham Surgery

Anesthesia was induced with 4% isoflurane. Once a surgical plane of anesthesia was confirmed via the loss of the pedal and corneal reflexes, isoflurane was maintained at 2% and ophthalmic ointment was applied to prevent eye desiccation during the procedure. Rats were positioned in the prone position on top of a sterile worksurface, complete with a thermal pad set to 37°C to minimize anesthesia-induced hypothermia. All burns were created unilaterally on the right hind paw. The right hind paw was aseptically prepared using alternating scrubs of Betadine and 70% ethanol. The full-thickness burn (i.e., “3^rd^ degree burn”) was created by pressing a 100° C soldering iron probe against the hind paw for 30 seconds. The slanted side of the soldering iron’s tip was used to create a horizontal burn across the plantar surface of the hind paw, perpendicular to the toes and heel. Sham rats underwent similar isoflurane anesthesia but did not receive a thermal injury. Immediately following surgery, burned rats received 1% silver sulfadiazine topical antibiotic cream directly on the burn injury. Silver sulfadiazine was administered to thermally injured rats once daily for the first four post-operative days.

### Spontaneous Pain Behavior – Gait Analysis

Spontaneous pain behaviors were captured using the CatWalk^TM^ XT (Noldus, Wageningen, Netherlands), an automatic gait analysis system that measures both dynamic and static gait parameters in rodents. Variables of interest included the pawprint area, pawprint width, minimum pawprint intensity, swing duration, swing speed, stand duration, and duty cycle. The CatWalk^TM^ XT apparatus^24^ comprises a 1.3m glass walking platform illuminated with a green backlight. Upon contact with the glass, the rat’s paw scatters the green light, illuminating any contact areas. The system is controlled by computer software (CatWalkXT, version 10.6) that captures the green pawprints relative to the light around the paw passing through the glass; the intensity of the illuminated pawprint area is proportional to the pressure exerted by the paw on the glass. A high-speed digital camera underneath the walking platform collects the live gait data and uploads the videos in real-time. Before baseline assessments were conducted, rats underwent 1 week of daily habituation sessions, wherein rats were allowed free exploration of the CatWalk^TM^ XT environment. Habituation sessions ended only when the rats reached the goalbox at the opposite end of the walkway. After each habituation or recording session, rats were returned to their home cages. During baseline and post-injury recordings, 10-15 gait videos were captured per rat to obtain at least three ‘compliant’ runs. ‘Compliant’ runs were trials in which the rat completed an unforced and uninterrupted walk along a predetermined area in the center of the platform (51.7cm in length). As such, recording sessions lasted approximately 10-15 minutes per rat, depending on how quickly a rat provided at least three compliant runs. The walkway was cleaned with a damp cloth between rats and with 70% ethanol between testing sessions.

### Evoked Pain Behavior – Mechanical Allodynia

Mechanical allodynia was assessed using a Dynamic Plantar Aesthesiometer (DPA; Ugo Basile, Italy). Rats were housed in non-restrictive Plexiglass enclosures situated above an elevated metal grid floor; the DPA sat underneath the metal grid floor, with its filament able to contact the rats’ hind paws. Rats were given 20 minutes to acclimate to the enclosure before testing began. The DPA uses a blunt, rigid 0.5mm metal filament on a movable force actuator to induce the paw withdrawal reflex and was set to ramp up pressure with linearly increasing force at a rate of 3 grams per second for 10 seconds, to a maximum of 30 grams of force. Each trial stopped when the paw was withdrawn from the filament or the cutoff time of 40 seconds was reached. The procedure was repeated for 3 trials per hind paw, with a 5-minute intertrial interval per paw. Despite the unilateral nature of the burn injury, all testing was conducted bilaterally, as the uninjured paw served as a within-subject control. All data, including the Paw Withdrawal Pressure (PWP), were collected automatically by the DPA’s onboard computer. The PWP was defined as the force in grams necessary to elicit the voluntary withdrawal of the paw. Mechanical allodynia testing was done after gait analysis, but before thermal hyperalgesia testing. The Plexiglass enclosure and metal grid floor were cleaned with a damp cloth between rats, and with 70% ethanol between sessions.

### Evoked Pain Behavior – Thermal Hyperalgesia

Thermal hyperalgesia was assessed using a plantar test for thermal stimulation (Ugo Basile, Italy), a movable infrared generator with an embedded paw withdrawal detector. Similar to mechanical allodynia testing, rats were housed in non-restrictive Plexiglass enclosures above a room temperature (∼30°C) glass floor plane. Thermal hyperalgesia testing occurred after mechanical allodynia testing, and rats received 20 minutes between these tests to acclimate to the thermal stimulation apparatus. The infrared beam of the thermal stimulator generated heat that traveled through the glass flooring and onto a focal point on the rat’s hind paw. The beam intensity was set to 80%, with a maximum trial length of 20 seconds to prevent thermal injury to the paw. Unlike the force intensity of the DPA, the thermal stimulator maintained a constant beam intensity during each trial. Three measurements were taken per hind paw for each thermal stimulation trial, with a 5-minute intertrial interval for each paw. Paw Withdrawal Latency (PWL) was the only behavioral metric collected. After thermal hyperalgesia testing, rats were returned to their home cages. The glass floor and Plexiglass enclosures were cleaned with a damp towel between rats, and with 70% ethanol between testing sessions.

### Alternative Pain Behavior (Anxiety) – Elevated Zero Maze (EZM)

The Elevated Zero Maze (Panlab) is a circular track divided into four quadrants, with a total diameter of 120 cm and a height of 62 cm off the floor. Of the four quadrants, two opposite quadrants are enclosed by darkened 31 cm-high acrylic walls. The other two quadrants are open to the room, with no walls along their edges. During trials, rats are allowed free exploration of the maze – there are no restraint devices, doors, or other impediments to movement about the maze’s circular track. At the beginning of each trial, the rat is placed in an open quadrant, with their head facing a closed quadrant. All rats were placed in the same open quadrant and faced the same closed quadrant for each EZM session. Using a Logitech camera mounted to the ceiling above the EZM, experimenters collected a single 10-minute video per rat per testing session; these videos were later analyzed using EthoVision (Noldus) tracking software. In the EthoVision software, experimenters analyzed two behavioral metrics: the number of entries into each quadrant and the cumulative duration of time spent in the open quadrants. The cumulative duration of time spent in an open area is a measure of anxiety in rodents.^25,26^ Additionally, the number of entries into a quadrant is a metric by which to examine anxiety-like and exploratory behavior in the maze. After each recording session, rats were returned to their home cages. Between rats, the EZM was cleaned with a damp cloth; between sessions, the EZM was cleaned with 70% ethanol.

### Humane Endpoints

To ensure the safety and well-being of our cohorts, we established several *a priori* humane endpoint criteria. Using the Pain Scale Table (see Supplementary Information, **Table S1**), experimenters monitored all rats daily for signs of excessive distress, including increased lethargy, weight loss of >20%, hunched posture, severe or prolonged porphyrin staining, or dehydration. Any rats that exhibited a morbid condition were removed from the study and humanely euthanized. While post-operative analgesia and systemic antibiotics were not offered due to their contraindication with experimental objectives, none of our rats exhibited signs of excess distress.

### Data Collection and Blinding

Rats were handled by experimenters of both sexes,^27^ and these same experimenters conducted the behavioral testing procedures and the daily monitoring care. Additionally, all behavioral tests were conducted during the light cycle; the majority of testing was conducted in the morning (7 AM-11 AM). Mechanical allodynia testing was always conducted before thermal hyperalgesia testing to avoid heat-induced numbing or heightened sensitivity affecting the accuracy of the mechanical allodynia data. Similarly, the CatWalk^TM^ XT session was always conducted prior to evoked pain testing to avoid the confounding effects of the evoked responses on spontaneous pain behavior. The EZM was conducted on separate days from the evoked and spontaneous testing to avoid confounding this sensitive psychological test. Due to the visual nature of the full-thickness burn injury, it was impossible to blind experimenters to group assignments during the experiment.

### Statistical Analyses

There were missing data for 6 total female rats (3 in the burned group, 3 in the control group) at the 28-, 42-, and 56-day timepoints. These values were considered missing at random and not due to their inherent nature. Missing data were handled using multiple imputations in SPSS (Version 31.0.0.0, IBM SPSS Software). We generated 5 imputed datasets to account for the ∼3% missing data. The imputation model included the outcome variable of interest, sex, injury group, and timepoint. For the three-way analyses of time, injury group, and sex, one of the 5 imputations were chosen at random using a random number generator. Three-way mixed ANOVAs were used to assess the effects of sex, injury group, and time on all evoked, spontaneous, and alternative pain variables. After a three-way mixed ANOVA, all variables were reanalyzed using sex-separated two-way mixed ANOVAs on the same imputation dataset. All data are mean ± SEM. Data quality for the three-way mixed ANOVAs was assessed using Shapiro-Wilk’s test for normal distribution, Levene’s Test of Equality of Error Variances for the assumption of homogenous error variances, and boxplots for outlier detection. Data quality for the two-way mixed ANOVAs was assessed using the same methods, but with the addition of Box’s Test for Equality of Covariances, and the examination of Studentized Residuals for further outlier detection. No outliers were removed from any variable. Where appropriate, significant pairwise comparisons of the ANOVAs are reported. For all statistical tests, when necessary and appropriate, variables were transformed to correct for non-normal distributions and heterogeneous error variances. All statistical analyses were conducted using SPSS; all graphs were created using GraphPad Prism (V.11.0.0, GraphPad Software, LLC.).

## Results

### Evoked Pain Responses: Male and Female Rats Display Convergent Reactions to Mechanical Stimulation, but Divergent Reactions to Noxious Thermal Stimulation

#### Mechanical Allodynia: Paw Withdrawal Pressure (PWP) of injured paw in grams (g)

For the pressure required to evoke withdrawal of the injured paw from a non-noxious stimulus, or Paw Withdrawal Pressure (PWP), the three-way interaction between time, injury group, and sex was not significant, *F*(11, 484) = 1.367, *p* = 0.185, partial η^2^ = 0.030. Rats with burn injuries showed a decrease in the PWP, over time, as indicated by a significant two-way interaction between time and injury group, *F*(11, 484) = 15.994, *p* < 0.001, partial η^2^ = 0.267, and significant main effects of time, *F*(11, 484) = 22.335, *p* < 0.001, partial η^2^ = 0.337, and injury group, *F*(1, 44) = 81.535, *p* < 0.001, partial η^2^ = 0.649. There was no significant two-way interaction between time and sex, *F*(11, 484) = 1.676, *p* = 0.076, partial η^2^ = 0.037, nor a significant main effect of sex, *F*(1, 44) = 0.819, *p* = 0.370, partial η^2^ = 0.018, indicating that there was a similar response for both males and females; however, the males showed a more exaggerated response to the burn injury as supported by a significant interaction between injury group and sex, *F*(1, 44) = 7.475, *p* = 0.009, partial η^2^ = 0.145. Because of these sex differences, two-way ANOVAs were conducted individually for males and females. For males (**Figure 2A**), there was a significant interaction between time and injury group, *F*(11, 242) = 11.995, *p* < 0.001, partial η^2^ = 0.353. The burn-injured group had a significantly lower PWP than controls at 1, 3, 7, 14, 21, 28, 42, 49, and 56 days post-burn, as supported by significant post hoc analyses. Females (**Figure 2B**) also had a significant interaction between time and injury group, *F*(11, 242) = 5.896, *p* < 0.001, partial η^2^ = 0.211. The burn-injured group had a significantly lower PWP than controls at 3, 7, 14, and 21 days post-burn, as supported by significant post hoc analyses.

**Figure 2.**
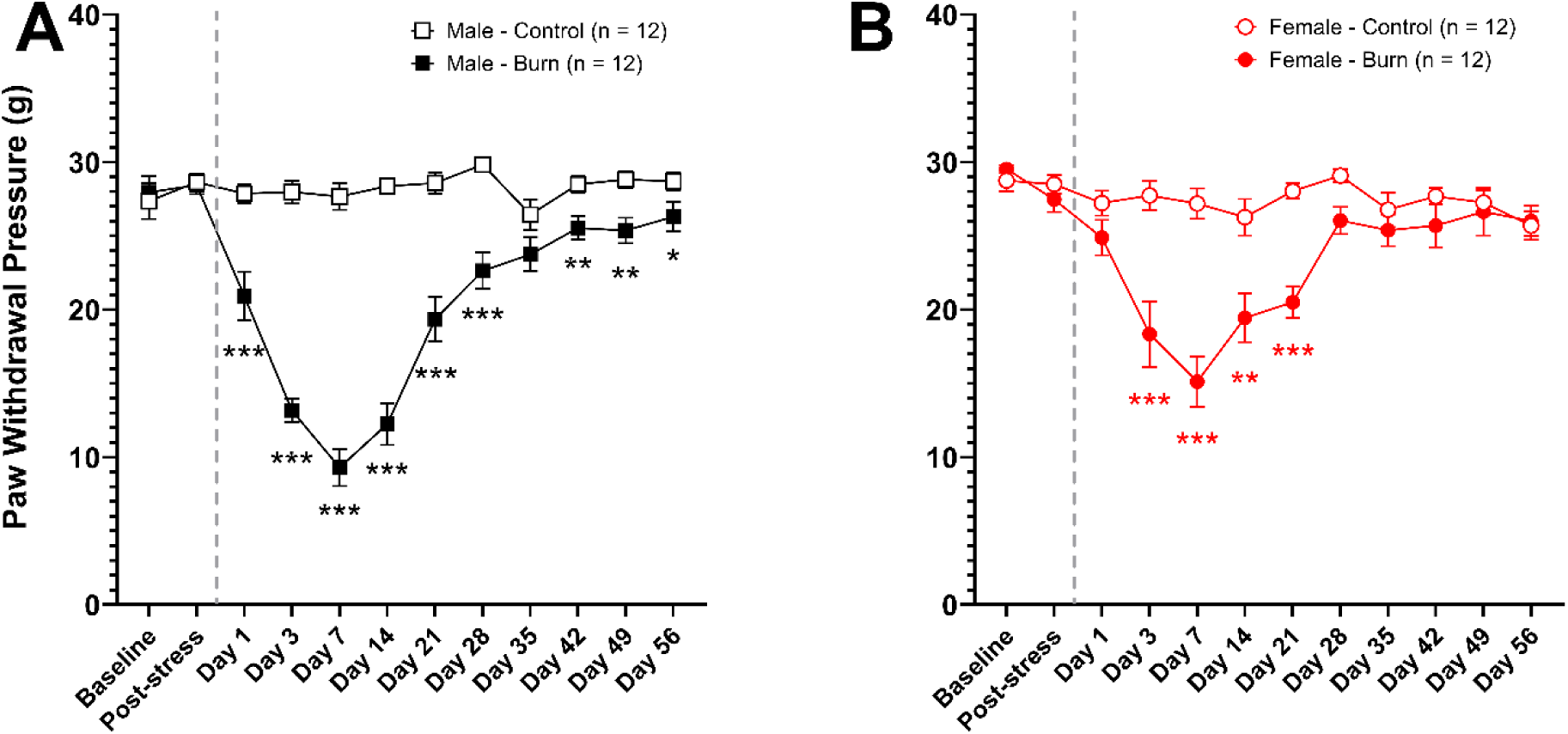
Evoked Pain Response: Mechanical Allodynia. Original, untransformed paw withdrawal pressure (PWP) in grams (g) displayed as mean ± SEM. **(A)** Male and **(B)** female rats demonstrated persistent mechanical allodynia (i.e., reduced PWP) over time following burn injury compared with controls. Males in the burn condition had a persistent post-burn reduction in PWP relative to control males, which persisted throughout the study period. Female rats in the burn condition also had reduced PWP post-burn relative to control females, but this allodynic reaction lasted only one month. The dashed line indicates the time of injury, with all time points thereafter indicating the number of days post-burn. \**p* < 0.05, \*\**p* < 0.01, \*\*\**p* < 0.001 indicates a significant difference between the Control and Burn conditions at individual timepoints.

#### Thermal Hyperalgesia: Paw Withdrawal Latency (PWL) Ratio (Injured/Uninjured Hind Paw)

Overall, the latency in seconds for the rat to withdraw their hind paw from a noxious thermal stimulus was significantly higher for females compared with males at all time points. Therefore, the paw withdrawal latency was analyzed as the ratio of the injured hind paw latency to the contralateral paw latency for each rat at each timepoint to normalize for this difference. The within-subject ratio was used for all further analyses. Results of the three-way repeated measures ANOVA found no significant three-way interaction between time, injury group, and sex, *F*(6.481, 285.171) = 1.792, *p* = 0.095, partial η^2^ = 0.039, ε = 0.589 (Greenhouse-Geisser). There were significant two-way interactions between time and injury group, *F*(6.481, 285.171) = 3.821, *p* < 0.001, partial η^2^ = 0.080, ε = 0.589 (Greenhouse-Geisser), time and sex, *F*(6.481, 285.171) = 2.105, *p* = 0.048, partial η^2^ = 0.046, ε = 0.589 (Greenhouse-Geisser), and injury group and sex, *F*(1, 44) = 18.808, *p* < 0.001, partial η^2^ = 0.299. There was also a significant main effect of time, *F*(6.481, 285.171) = 3.068, *p* = 0.005, partial η^2^ = 0.065, ε = 0.589 (Greenhouse-Geisser), and of sex, *F*(1, 44) = 13.902, *p* < 0.001, partial η^2^ = 0.240, but not of injury group, *F*(1, 44) = 0.302, *p* = 0.585, partial η^2^ = 0.007. Because there was a significant main effect of sex, the data were reanalyzed using separate two-way ANOVAs for each sex. For males (**Figure 3A**), there was a significant interaction between time and injury group, *F*(11, 242) = 2.943, *p* = 0.001, partial η^2^ = 0.118. Post hoc analyses indicated that the burn-injured group had increased sensitivity to the noxious thermal stimulus, as evidenced by a significantly lower PWL ratio compared with controls at 3, 7, 14, and 42 days post-burn. As anticipated, males displayed thermal hyperalgesia post-burn compared to uninjured, unstressed rats. Females (**Figure 3B**) showed a near-significant interaction between time and injury group, *F*(4.554, 100.194) = 2.353, *p* = 0.051, partial η^2^ = 0.097, ε = 0.414 (Greenhouse-Geisser). There was, however, a significant main effect of time, *F*(4.554, 100.194) = 2.636, *p* = 0.032, partial η^2^ = 0.107, ε = 0.414 (Greenhouse-Geisser), and a significant main effect of group, *F*(1,22) = 7.650, *p* = 0.011, partial η^2^ = 0.258. While there was a significant difference between the burn and control groups, the direction of the change in sensitivity was opposite that of males. Instead, females in the burn-injured group had a reduced sensitivity to the noxious thermal stimulus, as evidenced by a significantly higher PWL ratio compared with controls. The effect was most dramatic immediately after the burn injury on day 1 post-burn, and remained elevated, reaching significance at 21, 28, 35, 42, and 49 days post-burn. The acute increase in PWL ratio – i.e., a temporary resilience to noxious thermal stimulation – which was only present in the female rats, is a novel and unexpected result after full-thickness burn injury.

**Figure 3.**
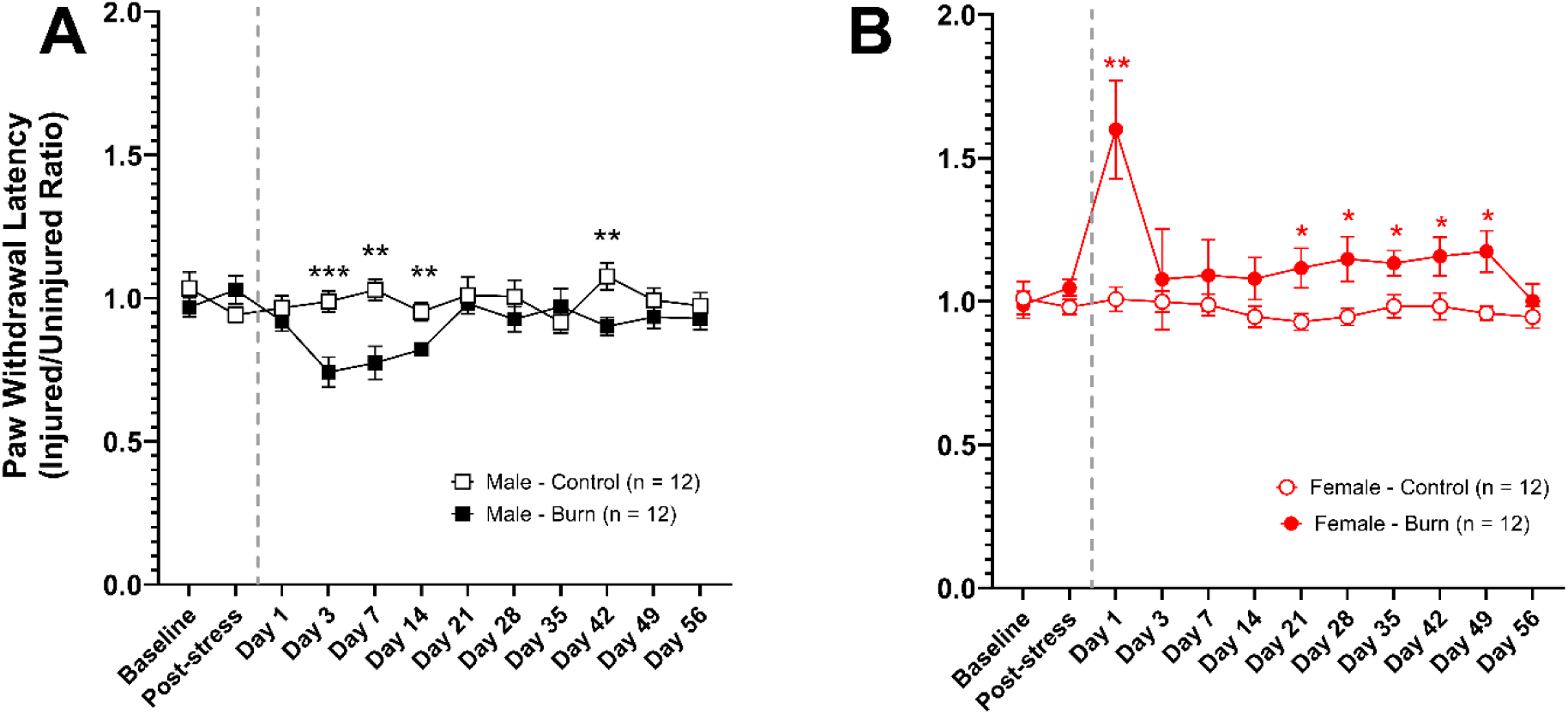
Evoked Pain Response: Thermal Hyperalgesia. Original, untransformed paw withdrawal latency (PWL) ratio displayed as mean ± SEM. **(A)** Male rats demonstrated thermal hyperalgesia (i.e., reduced PWL ratio) over time following burn injury compared with controls. **(B)** Female rats showed hypoalgesia over time following burn injury compared to controls. This effect peaked immediately following the injury but remained elevated until 56 days post-burn. The dashed line indicates the time of injury, with all time points thereafter indicating the number of days post-burn. \**p* < 0.05, \*\**p* < 0.01, \*\*\**p* < 0.001 indicates a significant difference between the Control and Burn conditions at individual timepoints.

### Gait Analysis – Static Features: Females with Burn Injuries Use a Smaller Portion of the Injured Foot with Each Step Compared with Sex-Matched Controls and Males with Burn Injuries

To account for differences in paw size between males and females, as well as potential changes over time as rats age, the variables used to assess gait are expressed as the ratio of the injured hind paw (right) to the uninjured hind paw (left) for each rat at each timepoint, thereby normalizing for this difference. The within-subject ratio was used for all further analyses.

#### Print Area Ratio (Injured/Uninjured Hind Paw)

The print area variable is the total surface area of the paw during the stance phase of a step and is used to assess weight-bearing and paw placement. Here, it is represented as a ratio of the injured paw surface area as a function of the surface area of the contralateral paw. A ratio of 1 means there is no difference between the two, while a decrease in the paw area ratio indicates that less weight is being applied to the injured paw. This is recognized as a sign of guarding. Results of the three-way repeated-measures ANOVA revealed no significant three-way interaction among time, injury group, and sex, *F*(11, 484) = 0.513, *p* = 0.895, partial η^2^ = 0.012. Overall, the rats with a burn injury had a reduced print area ratio, meaning that they walked on a smaller portion of their burned foot (right) than did uninjured controls, and this effect changed over time, as indicated by a significant two-way interaction between time and injury group, *F*(11, 484) = 4.261, *p* < 0.001, partial η^2^ = 0.088, and significant main effects of injury group, *F*(1, 44) = 31.508, *p* < 0.001, partial η^2^ = 0.417, and time, *F*(11, 484) = 4.306, *p* < 0.001, partial η^2^ = 0.089. A significant interaction between injury group and sex, *F*(1, 44) = 4.362, *p* = 0.043, partial η^2^ = 0.090, and a significant main effect of sex, *F*(1, 44) = 8.643, *p* = 0.005, partial η^2^ = 0.164, indicate that females had an exaggerated response to burn injury compared with males. Because there was a significant main effect of sex, the data were reanalyzed using separate two-way ANOVAs for males and females. For males (**Figure 4A**), there was a significant interaction between time and injury group, *F*(11, 242) = 2.931, *p* = 0.001, partial η^2^ = 0.118. Males with burn injuries had a temporary decrease in print area ratio compared to sex-matched controls that reached significance at 3, 7, 21, and 42 days post-burn. After a full-thickness burn, males stepped on their injured (right) hind paw with a smaller area than their uninjured (left) hind paw compared to healthy males, most severely in the first week post-burn. Females (**Figure 4B**) also showed a significant interaction between time and injury group, *F*(11, 242) = 2.401, *p* = 0.008, partial η^2^ = 0.098; however, their response was prolonged and there was a significant difference between injured and uninjured rats at 3, 7, 14, 28, 35, 42, 49, and 56 days post-burn. After a full-thickness burn, females demonstrated a persistent and prolonged reduction in the injured (right) hind paw print area relative to the contralateral healthy hind paw (left) area. The female response of using less of the injured hind paw with each step persisted longer than the equivalent male response.

**Figure 4.**
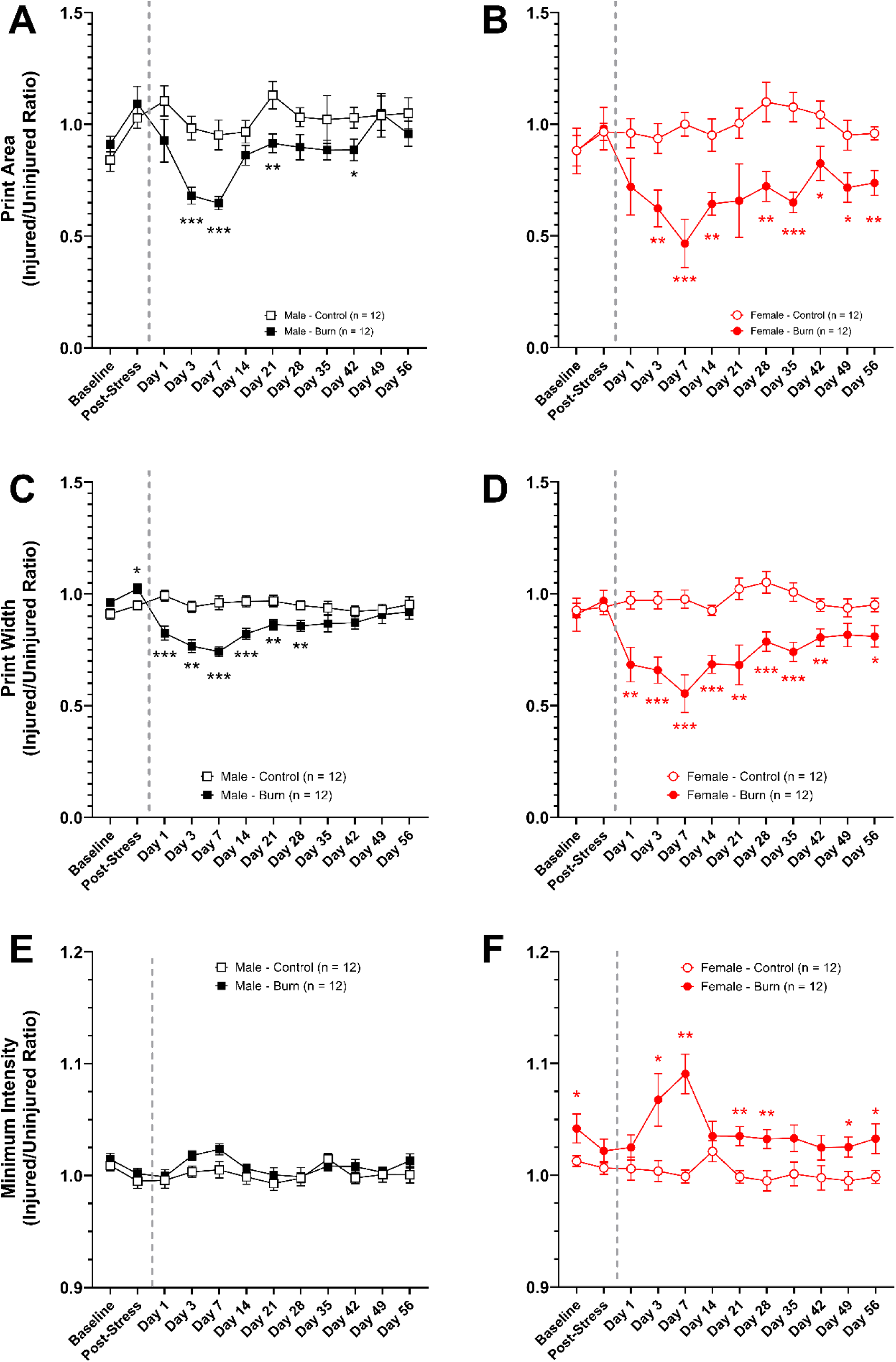
Static Gait Features: Paw Print Area Ratio. Original, untransformed paw print area ratio displayed as mean ± SEM. Male **(A)** and female **(B)** rats with a burn injury had a reduced print area ratio compared with sex-matched controls; for males, this was reflected in a temporary decrease; for females, it persisted throughout the post-burn period. Females with burn injuries also had significantly reduced print area ratio compared to males. **Paw Print Width Ratio.** Original, untransformed paw print width ratio displayed as mean ± SEM. Male **(C)** and female **(D)** rats with burn injury had reduced print width ratios compared with sex-matched controls. This effect was more pronounced and prolonged in females than in males. **Minimum Intensity Ratio.** Original, untransformed minimum paw intensity displayed as mean ± SEM. Male **(E)** rats did not show a significant change in minimum intensity ratio following burn injury, whereas female **(F)** rats showed a persistently increased minimum intensity ratio compared to sex-matched controls. This indicates that female rats are exerting greater pressure on a smaller area following burn injury, suggesting greater discomfort than males. These comparisons suggest that females put more pressure on their burned (right) paw than males. Taken together, females display an inability or unwillingness to use their entire right paw during post-burn ambulation compared to males. Additionally, females step more heavily on the small area of their burned paw that they use to ambulate and balance compared with males following burn injury. The dashed line indicates the time of injury, with all time points thereafter indicating the number of days post-burn. \**p* < 0.05, \*\**p* < 0.01, \*\*\**p* < 0.001 indicates a significant difference between the Control and Burn conditions at individual timepoints.

#### Print Width Ratio (Injured/Uninjured Hind Paw)

The print width variable is the width (or spread) of the paw during each step. As with print area, it is represented as a ratio of the injured paw width as a function of the width of the contralateral paw. A ratio of 1 indicates no difference between the two, while a decrease in the paw width ratio suggests the rat has a reduced spread of the injured paw when walking, suggesting increased guarding behaviors. Results of the three-way repeated measures ANOVA found no significant three-way interaction between time, injury group, and sex, *F*(6.960, 306.257) = 0.786, *p* = 0.599, partial η^2^ = 0.018, ε = 0.633 (Greenhouse-Geisser). As with the paw print area ratio, the rats with a burn injury had a reduced paw print width ratio, meaning that they put less weight on the injured paw, resulting in less paw spread during ambulation over time, as indicated by a significant two-way interaction between time and injury group, *F*(6.960, 306.257) = 11.087, *p* < 0.001, partial η^2^ = 0.201, and significant main effects of injury group, *F*(1, 44) = 33.256, *p* < 0.001, partial η^2^ = 0.430, and time, *F*(6.960, 306.257) = 6.741, *p* < 0.001, partial η^2^ = 0.133. Similar to the paw print area ratio, the width ratio showed an exaggerated response in female rats. This was supported by a significant interaction between injury group and sex, *F*(1, 44) = 6.816, *p* = 0.012, partial η^2^ = 0.134; however, the main effect of sex did not reach significance, *F*(1, 44) = 3.207, *p* = 0.080, partial η^2^ = 0.068. To further investigate the significant interaction between sex and injury group, the data were reanalyzed using separate two-way ANOVAs for males and females. For males (**Figure 4C**), there was a significant interaction between time and injury group, *F*(11, 242) = 8.076, *p* < 0.001, partial η^2^ = 0.269. Males with burn injuries had a decrease in print width ratio compared to sex-matched controls that reached significance at the Post-Stress timepoint, and 1, 3, 7, 14, 21, and 28 days post-burn. Similarly, females (**Figure 4D**) also showed a significant interaction between time and injury group, *F*(5.902, 129.8) = 5.303, p < 0.001, partial η^2^ = 0.194, ε = 0.5366 (Greenhouse-Geisser); however, their response was prolonged, with a significant difference between injured and uninjured rats at 1, 3, 7, 14, 21, 28, 35, 42, and 56 days post-burn. The reduction in print width ratio, compared to sex-matched healthy controls, persisted for weeks longer in burned females than in burned males. Additionally, the magnitude of the reduction in the print width ratio was larger in burned females than in burned males. Taken with the results for print area, these data further indicate that burned rats used a smaller portion of the burned foot in each step than the unburned foot, but that female rats demonstrated a more severe and persistent gait alteration.

#### Minimum Intensity Ratio (Injured/Uninjured Hind Paw)

The minimum intensity variable represents the minimum force applied to the paw during each step. Here, it is represented as a ratio of the minimum intensity of the injured paw (right) as a function of that of the contralateral paw (left). A ratio of 1 means there is no difference between the two, while an increase in the minimum intensity ratio indicates that the lowest force applied during each step is higher, which suggests gait dysfunction. Results of the three-way repeated measures ANOVA found no significant three-way interaction between time, injury group, and sex, *F*(7.836, 344.798) = 1.136, *p* = 0.339, partial η^2^ = 0.025, ε = 0.712 (Greenhouse-Geisser). The minimum intensity ratio increased over time following burn injury, as indicated by a significant two-way interaction between time and injury group, *F*(7.836, 344.798) = 2.319, *p* = 0.020, partial η^2^ = 0.050, and significant main effects of injury group, *F*(1, 44) = 27.556, *p* < 0.001, partial η^2^ = 0.385, and time, *F*(7.836, 344.798) = 3.538, *p* < 0.001, partial η^2^ = 0.074. As with the other static gait measures, the minimum intensity ratio showed an exaggerated response in female rats. This was supported by a significant interaction between injury group and sex, *F*(1, 44) = 11.901, *p* = 0.001, partial η^2^ = 0.213, and a significant main effect of sex, *F*(1, 44) = 15.020, *p* < 0.001, partial η^2^ = 0.254. To further investigate the significant interaction between sex and injury group, the data were reanalyzed using separate two-way ANOVAs for males and females. There was no significant interaction between time and injury group for males (**Figure 4E**), *F*(11, 242) = 0.733, *p* = 0.706, partial η^2^ = 0.032, nor was there a significant main effect of group, *F*(1, 22) = 3.086, *p* = 0.093, partial η^2^ = 0.123. There was a significant main effect of time, *F*(11, 242) = 2.775, *p* = 0.002, partial η^2^ = 0.112. Overall, the minimum intensity ratio was not significantly altered by burn injury in the male rats, although there was some fluctuation over time. Conversely, there was a significant interaction between time and injury group for the females (**Figure 4F**), *F*(11, 242) = 2.046, *p* = 0.025, partial η^2^ = 0.085, with differences between the burn and control groups reaching significance at the pre-stress baseline, and 3, 7, 21, 28, 49, and 56 days post-burn. The differences between female groups at the pre-stress baseline were smaller than those seen post-burn. As with the variability in males, this may be due to the variability in this measure. It is unlikely to be indicative of substantive preexisting differences between the groups, given that the pre-injury timepoint (Post-Stress) did not show a significant group difference. As with the other static gait measures, the minimum intensity ratio showed an exaggerated response in female rats following burn injury.

### Gait Analysis – Dynamic Features: Females with Burn Injuries Display a Pronounced, Prolonged Disturbance of Dynamic Gait Features Compared to Males with Burn Injury and Sex-matched Controls

#### Swing Ratio (Injured/Uninjured Hind Paw)

The swing variable is the amount of time the paw is in the air during a step cycle. A longer swing time is indicative of guarding behavior. As with the other gait measures, swing is represented as a ratio of the time the injured paw (right) is in the air as a function of the time the uninjured contralateral paw (left) is in the air. A ratio of 1 means there is no difference between the two, while an increase in the swing ratio indicates that the injured paw is held in the air for a longer period of time. Results of the three-way repeated measures ANOVA found no significant three-way interaction between time, injury group, and sex, *F*(3.532, 155.389) = 1.526, *p* = 0.203, partial η^2^ = 0.034, ε = 0.321 (Greenhouse-Geisser). The swing ratio increased over time following burn injury, as indicated by a significant two-way interaction between time and injury group, *F*(3.532, 155.389) = 8.003, *p* < 0.001, partial η^2^ = 0.154, ε = 0.321 (Greenhouse-Geisser), and significant main effects of injury group, *F*(1, 44) = 13.042, *p* < 0.001, partial η^2^ = 0.229, and time, *F*(3.532, 155.389) = 6.883, *p* < 0.001, partial η^2^ = 0.135, ε = 0.321 (Greenhouse-Geisser). As with the static gait measures, the swing ratio showed an exaggerated response in female rats, as indicated by a significant main effect of sex, *F*(1, 44) = 5.859, *p* = 0.020, partial η^2^ = 0.118. There was no significant interaction between injury group and sex, *F*(1, 44) = 2.812, *p* = 0.101, partial η^2^ = 0.060. To further investigate the significant main effect of sex, the data were reanalyzed using separate two-way ANOVAs for males and females. For males (**Figure 5A**), there was a significant interaction between time and injury group, *F*(6.026, 132.6) = 4.154, *p* < 0.001, partial η^2^ = 0.159, ε = 0.5478 (Greenhouse-Geisser). Post hoc analyses indicated that the burn-injured group had increased swing time, as evidenced by a significantly higher swing ratio compared with controls at 3, 7, and 28 days post-burn. Females (**Figure 5B**) also showed a significant interaction between time and injury group, *F*(3.236, 71.201) = 5.232, *p* = 0.002, partial η^2^ = 0.192, ε = 0.294 (Greenhouse-Geisser), with the burn group exhibiting increased swing ratios compared with the controls at 1, 3, 7, 21, 28, 49, and 56 days post-burn. In summary, rats in the burn condition kept the injured foot in the air longer, step-to-step, than the uninjured foot; this response was exaggerated in females, persisting throughout the entire post-burn period, but resolved in males within one month.

**Figure 5.**
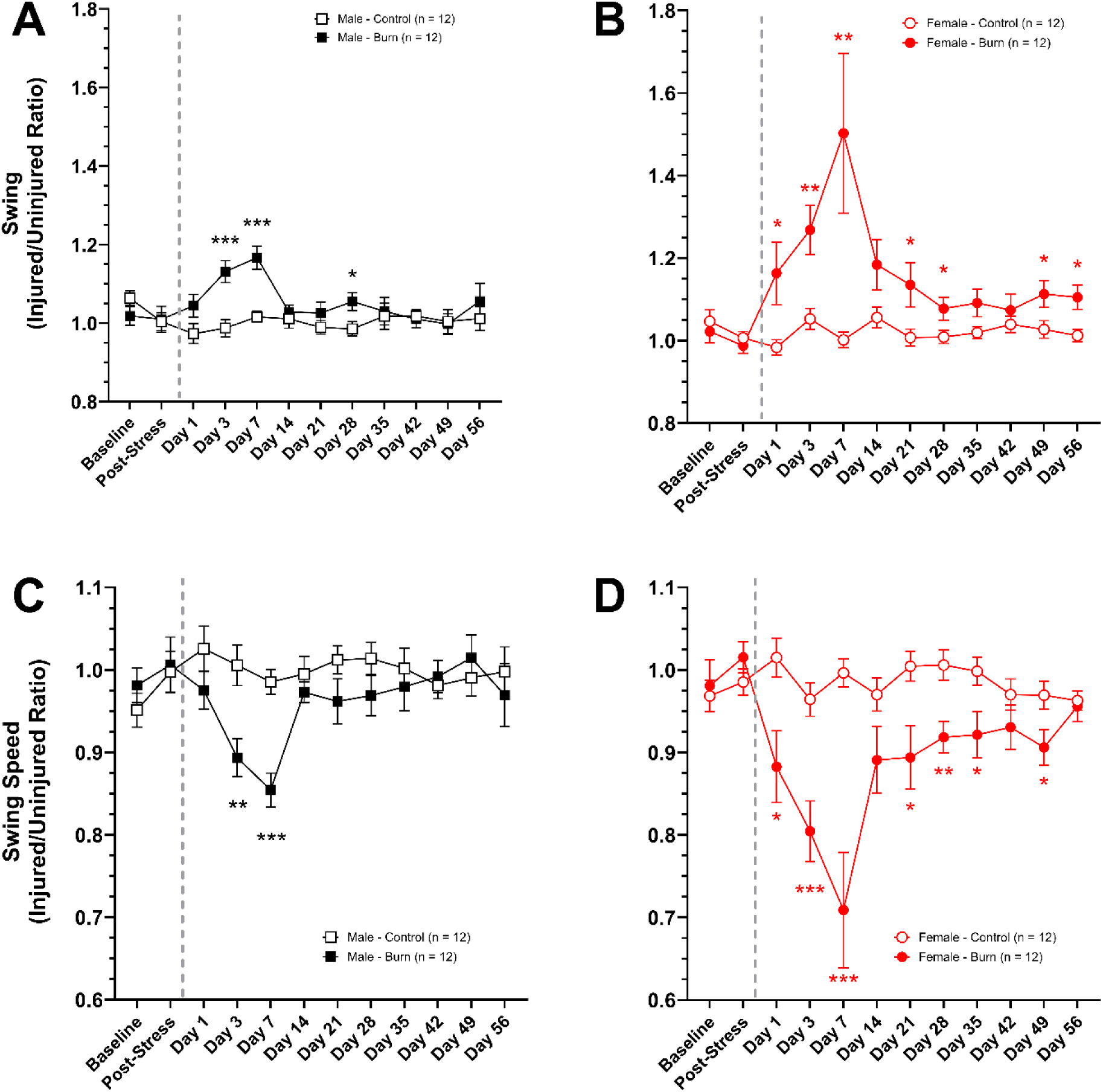
Dynamic Gait Features: Swing Ratio. Original, untransformed swing ratio displayed as mean ± SEM. Male **(A)** and female **(B)** rats had an increased swing ratio following burn injury, indicating that the burned (right) foot spent longer in the air between steps than the right feet of the control groups. Females had a more pronounced change in swing ratio within the first month post-burn compared with the males. **Swing Speed Ratio.** Original, untransformed swing speed ratio displayed as mean ± SEM. Male **(C)** and female **(D)** rats with burn injury had reduced swing speed ratios compared with sex-matched controls, indicating that their burned (right) foot displaced a smaller distance every second of step-to-step motion compared to the control groups. This effect was more pronounced and prolonged in females than in males. The dashed line indicates the time of injury, with all time points thereafter indicating the number of days post-burn. \**p* < 0.05, \*\**p* < 0.01, \*\*\**p* < 0.001 indicates a significant difference between the Control and Burn conditions at individual timepoints.

#### Swing Speed Ratio (Injured/Uninjured Hind Paw)

The swing speed variable is the speed of the paw during a stride. A slower swing speed is indicative of pain or guarding behaviors. Swing speed is represented as a ratio of the injured paw (right) as a function of the speed of the uninjured contralateral paw (left). A ratio of 1 means there is no difference between the two, while a decrease in the swing speed ratio indicates that the injured paw is being moved at a slower rate than the contralateral paw. Results of the three-way repeated measures ANOVA found no significant three-way interaction between time, injury group, and sex, *F*(5.409, 237.977) = 1.595, *p* = 0.157, partial η^2^ = 0.035, ε = 0.492 (Greenhouse-Geisser). The swing speed ratio decreased over time following burn injury, as indicated by a significant two-way interaction between time and injury group, *F*(5.409, 237.977) = 10.016, *p* < 0.001, partial η^2^ = 0.185, ε = 0.492 (Greenhouse-Geisser), and significant main effects of injury group, *F*(1, 44) = 12.868, *p* < 0.001, partial η^2^ = 0.226, and time, *F*(5.409, 237.977) = 8.626, *p* <0.001, partial η^2^ = 0.164, ε = 0.492 (Greenhouse-Geisser). There was an exaggerated response in female rats, as indicated by a significant main effect of sex, *F*(1, 44) = 5.477, *p* = 0.024, partial η^2^ = 0.111. There was no significant interaction between injury group and sex, *F*(1, 44) = 2.768, *p* = 0.103, partial η^2^ = 0.059. To further investigate the significant main effect of sex, the data were reanalyzed using separate two-way ANOVAs for males and females. For males (**Figure 5C**), there was a significant interaction between time and injury group, *F*(11, 242) = 3.684, *p* < 0.001, partial η^2^ = 0.143. Post hoc analyses indicated that the burn-injured group had a slower swing speed, as evidenced by a significantly lower swing speed ratio compared with controls at 3- and 7-days post-burn. Females (**Figure 5D**) also showed a significant interaction between time and injury group, *F*(4.288, 94.34) = 6.470, p < 0.001, partial η^2^ = 0.227, ε = 0.3898 (Greenhouse-Geisser). Post hoc tests indicate that females in the burn-injured group had a reduced swing speed overall, as evidenced by a significantly lower swing speed ratio. This reached significance at 1, 3, 7, 21, 28, 35, and 49 days post-burn. After burn injury, female rats had a slower swing speed for the injured foot than the uninjured foot. While male burned rats also displayed a slowed swing speed compared to healthy males, their presentation was mild and acute in comparison to burned females.

#### Stand Ratio (Injured/Uninjured Hind Paw)

The stand variable is the time the paw is in contact with the glass surface during a step cycle. Rats with injuries tend to have reduced contact time of the injured paw with the glass surface. Here, it is represented as a ratio of the injured paw as a function of the contralateral paw. A ratio of 1 means there is no difference between the two, while a decrease in the stand ratio indicates the paw is in contact with the glass for a shorter duration during each step. Results of the three-way repeated measures ANOVA found no significant three-way interaction between time, injury group, and sex, *F*(5.594, 246.116) = 1.117, *p* = 0.353, partial η^2^ = 0.025, ε = 0.509 (Greenhouse-Geisser). The stand ratio decreased over time following burn injury, as indicated by a significant two-way interaction between time and injury group, *F*(5.594, 246.116) = 6.481, *p* < 0.001, partial η^2^ = 0.128, ε = 0.509 (Greenhouse-Geisser), and significant main effects of injury group, *F*(1, 44) = 12.293, *p* = 0.001, partial η^2^ = 0.218, and time, *F*(5.594, 246.116) = 5.381, *p* < 0.001, partial η^2^ = 0.109, ε = 0.509 (Greenhouse-Geisser). There was an exaggerated response in female rats, as indicated by a significant main effect of sex, *F*(1, 44) = 11.426, *p* = 0.002, partial η^2^ = 0.206. There was no significant interaction between injury group and sex, *F*(1, 44) = 2.985, *p* = 0.091, partial η^2^ = 0.064. To further investigate the significant main effect of sex, the data were reanalyzed using separate two-way ANOVAs for males and females. For males (**Figure 6A**), there was a significant interaction between time and injury group, *F*(11, 242) = 3.628, *p* < 0.001, partial η^2^ = 0.142. Post hoc analyses indicated that the burn-injured group had a lower stand ratio than controls at 3, 7, and 14 days post-burn, and a higher stand ratio at 42 days post-burn. Females (**Figure 6B**) also showed a significant interaction between time and injury group, *F*(4.336, 95.39) = 3.847, p = 0.0049, partial η^2^ = 0.149, ε = 0.3942 (Greenhouse-Geisser). Post hoc tests indicate that females in the burn-injured group had a reduced stand ratio that reached significance compared to controls at 3, 7, 14, 28, and 35 days post-burn. After burn injury, female rats contacted the glass surface for less time on the burned hind paw than on the uninjured hind paw. Males also reduced the contact time on the burned paw compared to healthy rats, but this change lasted only two weeks post-burn, compared to one month in the females.

**Figure 6.**
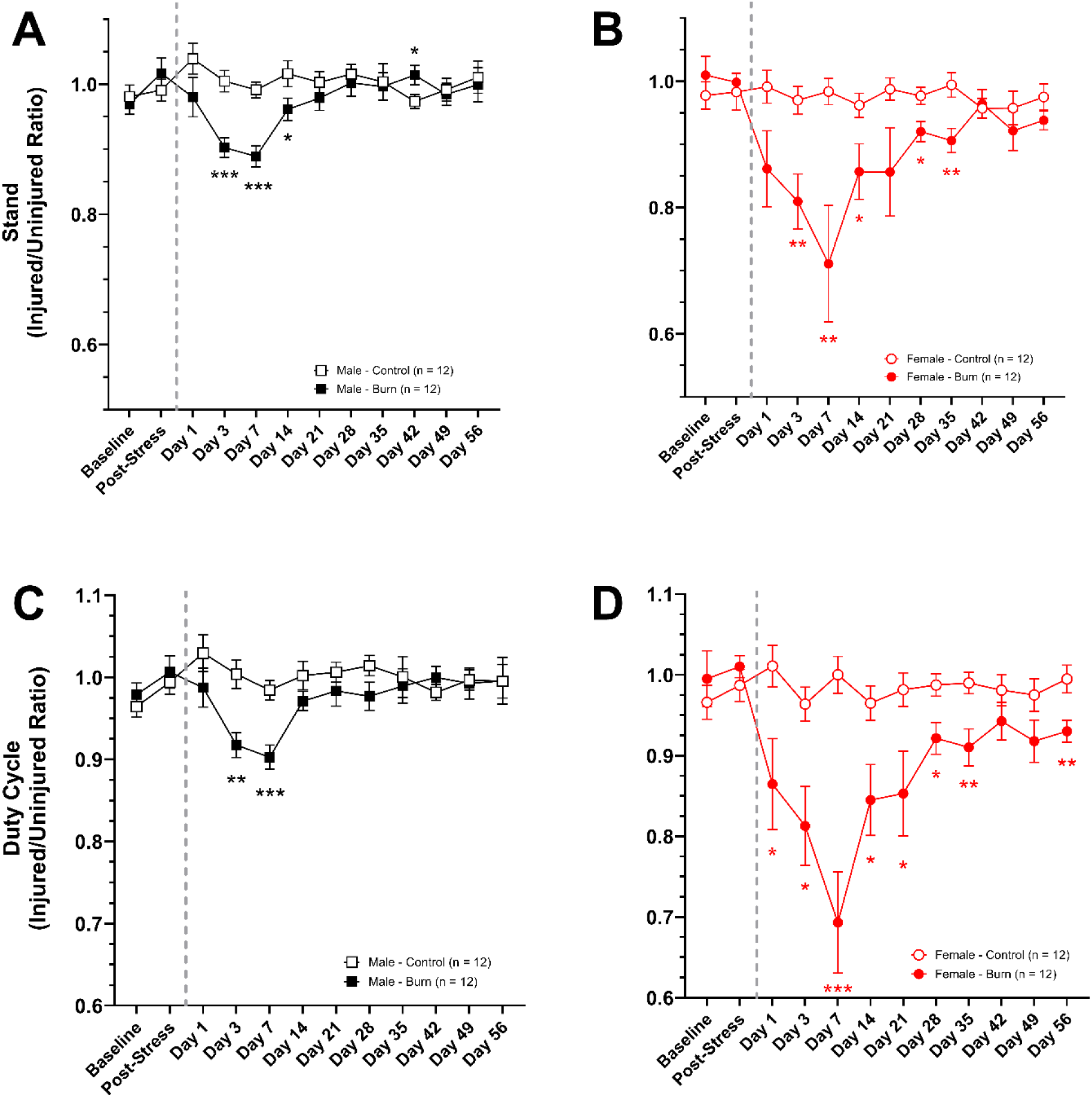
Dynamic Gait Features: Stand Ratio. Original, untransformed stand ratio displayed as mean ± SEM. Male **(A)** and female **(B)** rats with a burn injury had a reduced stand ratio compared with sex-matched controls; for males, this was reflected in a temporary (< 2 week) decrease; for females, it persisted throughout the post-burn period. Females with burn injuries also had significantly reduced stand ratios compared to males. **Duty Cycle Ratio.** Original, untransformed duty cycle ratio displayed as mean ± SEM. Male **(C)** and female **(D)** rats with burn injury had reduced duty cycle ratios than sex-matched controls. Females with burn injuries had a more pronounced effect compared with males throughout the post-burn period. The dashed line indicates the time of injury, with all time points thereafter indicating the number of days post-burn. \**p* < 0.05, \*\**p* < 0.01, \*\*\**p* < 0.001 indicates a significant difference between the Control and Burn conditions at individual timepoints.

#### Duty Cycle Ratio (Injured/Uninjured Hind Paw)

Duty cycle is the percentage of a full step cycle during which the paw is in contact with the glass surface, relative to the total duration of a complete step cycle. A reduction in duty cycle is indicative of pain or discomfort. As with the other gait measures, the duty cycle is represented as a ratio of the injured paw (right) to the uninjured contralateral paw (left), with a duty cycle ratio of 1 indicating no difference between the two paws and a decreased duty cycle ratio indicating more pain or discomfort in the injured paw. Results of the three-way repeated measures ANOVA found no significant three-way interaction between time, injury group, and sex, *F*(6.372, 280.378) = 2.211, *p* = 0.039, partial η^2^ = 0.048, ε = 0.579 (Greenhouse-Geisser). The duty cycle ratio decreased over time following burn injury, as indicated by a significant two-way interaction between time and injury group, *F*(6.372, 280.378) = 8.325, *p* < 0.001, partial η^2^ = 0.159, ε = 0.579 (Greenhouse-Geisser), and significant main effects of injury group, *F*(1, 44) = 14.816, *p* < 0.001, partial η^2^ = 0.252, and time, *F*(6.372, 280.378) = 7.239, *p* < 0.001, partial η^2^ = 0.141, ε = 0.579 (Greenhouse-Geisser). There was an exaggerated response in female rats, as indicated by a significant two-way interaction between sex and injury group, *F*(1, 44) = 5.451, *p* = 0.024, partial η^2^ = 0.110, as well as a significant main effect of sex, *F*(1, 44) = 10.812, *p* = 0.002, partial η^2^ = 0.197. To further investigate the significant interaction and main effect of sex, the data were reanalyzed using separate two-way ANOVAs for males and females. For males (**Figure 6C**), there was a significant interaction between time and injury group, *F*(5.699, 125.4) = 3.181, *p* = 0.0070, partial η^2^ = 0.126, ε = 0.5181 (Greenhouse-Geisser). Post hoc analyses indicated that the burn-injured group had a lower duty cycle ratio than controls at 3- and 7-days post-burn. Females (**Figure 6D**) also showed a significant interaction between time and injury group, *F*(5.209, 114.6) = 5.856, p < 0.001, partial η^2^ = 0.210, ε = 0.4735 (Greenhouse-Geisser). Post hoc tests indicate that females in the burn-injured group had a lower duty cycle ratio that reached significance compared with the control group at 1, 3, 7, 14, 21, 28, 35, and 56 days post-burn. Overall, following burn injury, injured rats had a reduced duty cycle ratio compared with controls, and this was most pronounced and persistent in females.

### Elevated Zero Maze: Males, but not Females, with Burn Injuries Demonstrate Post-Burn Anxiety-like Behavior

#### Time Spent in the Open Areas

The total time spent in the open areas of the elevated zero maze (EZM) is a sensitive measurement of rodent anxiety. Anxious rodents will not venture into the open, uncovered spaces of the EZM, but will instead stay in the walled “closed” sections of the maze. The three-way interaction of time, injury group, and sex was not significant, *F*(3.067, 134.936) = 0.495, *p* = 0.690, partial η^2^ = 0.011, ε = 0.767 (Greenhouse-Geisser). The two-way interactions between time and sex, *F*(3.067, 134.936) = 0.858, *p* = 0.467, partial η^2^ = 0.019, ε = 0.767 (Greenhouse-Geisser), and time and injury group, *F*(3.067, 134.936) = 1.541, *p* = 0.206, partial η^2^ = 0.034, ε = 0.767 (Greenhouse-Geisser), were not significant; however, the two-way interaction between injury group and sex was nearing significance, *F*(1, 44) = 3.978, *p* = 0.052, partial η^2^ = 0.083. There was also a significant main effect of time, *F*(3.067, 134.936) = 8.218, *p* < 0.001, partial η^2^ = 0.157, ε = 0.767 (Greenhouse-Geisser), and injury group, *F*(1, 44) = 5.014, *p* = 0.030, partial η^2^ = 0.102, but no significant main effect of sex, *F*(1, 44) = 1.416, *p* = 0.240, partial η^2^ = 0.031. The data were reanalyzed using separate two-way ANOVAs for males and females. For males (**Figure 7A**), there was not a significant interaction between time and injury group, *F*(4, 88) = 1.792, *p* = 0.137, partial η^2^ = 0.075; however, there was a significant main effect of injury group, *F*(1, 22) = 7.830, *p* = 0.010, partial η^2^ = 0.262, indicating that the burn group spent less time in the open areas compared with controls at all post-burn timepoints. There was also a significant main effect of time, *F*(4, 88) = 3.785, *p* = 0.007, partial η^2^ = 0.147, suggesting that over the study period, healthy males spent increasing time in the open areas of the EZM as they became familiar with the maze; burned males, conversely, did not substantially increase their time in the open areas, suggesting greater anxiety than the control males despite also becoming more familiar with the EZM. Females (**Figure 7B**) also failed to show a significant interaction between time and injury group, *F*(2.816, 61.959) = 0.280, p = 0.828, partial η^2^ = 0.013, ε = 0.704 (Greenhouse-Geisser); however, there was a significant main effect of time, *F*(2.816, 61.959) = 5.255, *p* = 0.003, partial η^2^ = 0.193, ε = 0.704 (Greenhouse-Geisser), which indicated the rats were spending more time in the open area as the study progressed. This was similar for both the control and burn rats, as indicated by a non-significant main effect of injury group, *F*(1, 22) = 0.035, *p* = 0.853, partial η^2^ = 0.002. This finding suggests that, unlike males, female rats in the burn condition did not express signs of post-burn anxiety.

**Figure 7.**
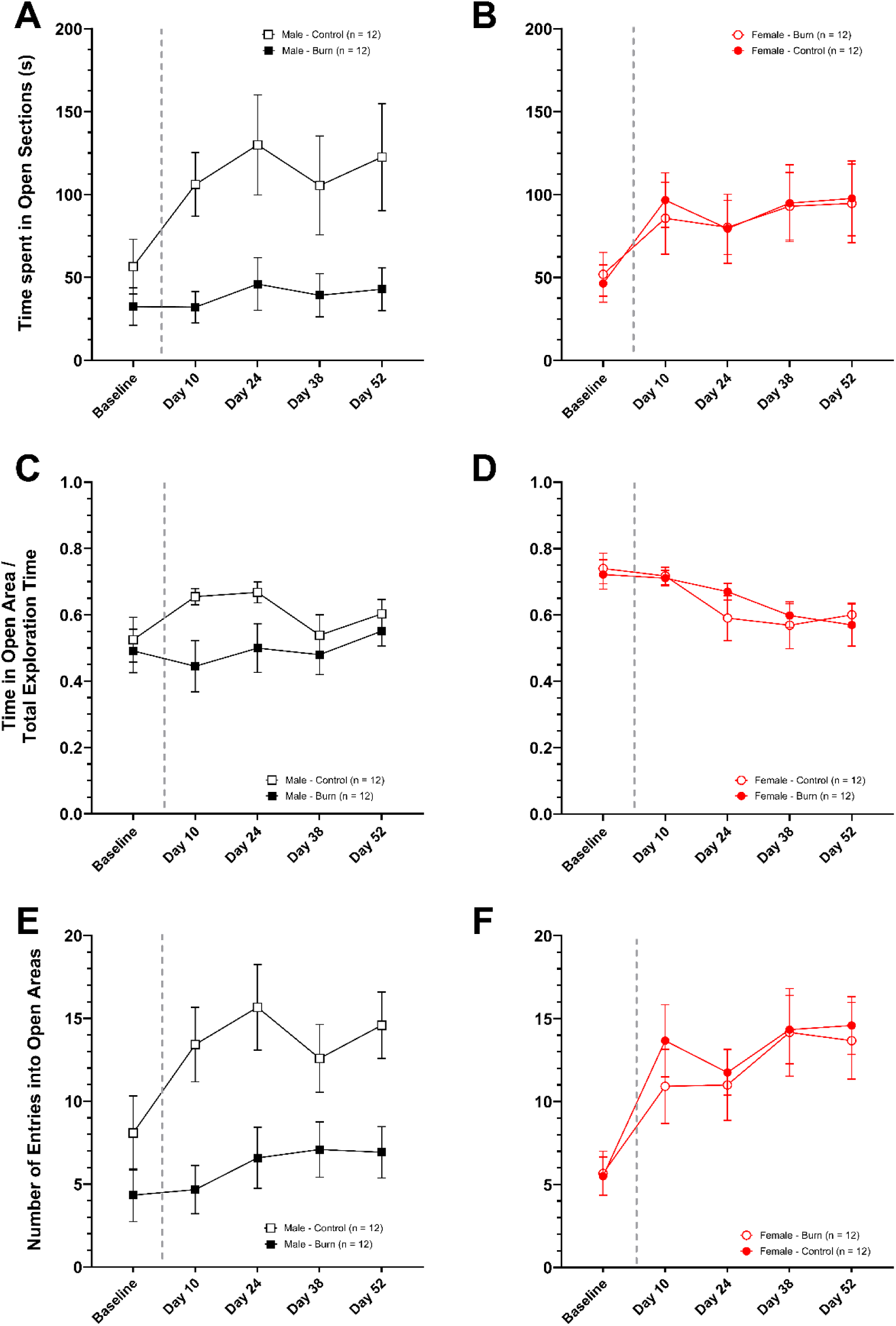
Elevated Zero Maze (EZM): Time in Open Areas. Time spent in the open areas of the elevated zero maze, in seconds, is presented as mean ± SEM. Male **(A)** rats with a burn injury spent less time in the open areas of the EZM following baseline, compared with controls. This effect was not observed in female **(B)** rats, which showed an increase in time spent in open areas after baseline that remained constant and did not differ based on burn condition. **Time in Open Area as a Function of Total Exploration.** The ratio of time spent in open areas to total exploration time was assessed to account for differences in activity between groups. Data are displayed as mean ± SEM. Male **(C)** rats with a burn injury spent less active time in the open areas of the EZM compared with controls. This effect was not observed in female **(D)** rats, which showed a general decrease in active time spent in open areas over time with no difference between burn and control conditions. **Entries into Open Areas.** The number of entries into the open area of the EZM is displayed as mean ± SEM. Males **(E)** with burn injuries entered the open area far fewer times than sex-matched controls at all post-burn timepoints. Females **(F)** with burn injuries did not differ significantly from their sex-matched controls but did enter the open area significantly more over time. Taken together, these data suggest that only the males with burn injuries had anxiety-like behavior, as they were the only group not to increase their open area exploration in the EZM over time. The dashed line indicates the time of injury, with all time points thereafter indicating the number of days post-burn.

#### Time Spent in the Open Area as a Function of Total Exploration Time

To ensure that the interpretation of time spent in the open area accurately reflected changes in anxiety post-burn, rather than burn-induced changes in locomotor activity or ability, the open area exploration data were reanalyzed as a function of total exploration time (i.e., total movement time) on the EZM. A lower ratio of open time to total time exploring indicates that rats spent a lower proportion of the exploratory time specifically exploring the open areas. Results of the three-way repeated-measures ANOVA showed no significant three-way interaction among time, injury group, and sex, *F*(3.107, 136.704) = 1.370, *p* = 0.254, partial η^2^ = 0.030, ε = 0.777 (Greenhouse-Geisser). Males and females responded differently over time following burn injury, as indicated by a significant two-way interaction between time and sex, *F*(3.107, 136.704) = 3.806, *p* = 0.011, partial η^2^ = 0.080, ε = 0.777 (Greenhouse-Geisser), and a significant main effect of sex, *F*(1, 44) = 9.439, *p* = 0.004, partial η^2^ = 0.177. There were no other significant interactions, including the two-way interactions between time and injury group, *F*(3.107, 136.704) = 0.648, *p* = 0.591, partial η^2^ = 0.015, ε = 0.777 (Greenhouse-Geisser), and the two-way interaction between sex and injury group, *F*(1, 44) = 2.943, *p* = 0.093, partial η^2^ = 0.063. There was no significant main effect of time, *F*(3.107, 136.704) = 2.272, *p* = 0.081, partial η^2^ = 0.049, ε = 0.777 (Greenhouse-Geisser), or injury group, *F*(1, 44) = 1.949, *p* = 0.170, partial η^2^ = 0.042. To further investigate the significant main effect of sex and the significant time-by-sex interaction, the data were reanalyzed using separate two-way ANOVAs for males and females. For males (**Figure 7C**), there was not a significant interaction between time and injury group, *F*(2.974, 65.433) = 1.061, *p* = 0.372, partial η^2^ = 0.046, ε = 0.744 (Greenhouse-Geisser), nor was there a significant main effect of time, *F*(2.974, 65.433) = 1.285, *p* = 0.287, partial η^2^ = 0.055, ε = 0.744 (Greenhouse-Geisser). The main effect of injury group approached significance, *F*(1, 22) = 3.983, *p* = 0.058, partial η^2^ = 0.153, suggesting that the burn group spent less time in open areas as a function of total exploration than the controls. Females (**Figure 7D**) also failed to show a significant interaction between time and injury group, *F*(4, 88) = 0.601, p = 0.663, partial η^2^ = 0.027; however, there was a significant main effect of time, *F*(4,88) = 5.966, *p* < 0.001, partial η^2^ = 0.213. There was a decrease in active time spent in the open area as a function of overall exploration over time, and this was similar for both the control and burn rats, as indicated by a non-significant main effect of injury group, *F*(1, 22) = 0.065, *p* = 0.801, partial η^2^ = 0.003. This suggests that, as the study progressed, all female rats reduced the amount of time spent in the open area of the maze relative to their total time spent moving in the EZM. While this could be indicative of increased anxiety over time, the increase in exploration time in the open areas for all females (**Figure 7B**) does not support this interpretation.

#### Entries to Open Area

The number of entries into the open areas of the EZM is another means to capture exploratory and anxiety-like behavior. Anxious rats will enter open areas less than non-anxious rats. Results of the three-way repeated-measures ANOVA showed no significant interaction among time, injury group, and sex, *F*(2.623, 115.395) = 0.376, *p* = 0.743, partial η^2^ = 0.008, ε = 0.656 (Greenhouse-Geisser). There was a significant main effect of time, *F*(2.623, 115.395) = 15.609, *p* < 0.001, partial η^2^ = 0.262, ε = 0.656 (Greenhouse-Geisser), and injury group, *F*(1, 44) = 6.329, *p* = 0.016, partial η^2^ = 0.126, but not a significant interaction between time and injury group, *F*(2.623, 115.395) = 1.425, *p* = 0.242, partial η^2^ = 0.031, ε = 0.656 (Greenhouse-Geisser). There was not a significant main effect of sex, *F*(1, 44) = 1.878, *p* = 0.178, partial η^2^ = 0.041; however, the interactions between time and sex, *F*(2.623, 115.395) = 2.678, *p* = 0.058, partial η^2^ = 0.057, ε = 0.656 (Greenhouse-Geisser), and between sex and injury group, *F*(1, 44) = 3.796, *p* = 0.058, partial η^2^ = 0.079, were nearing significance. The data were reanalyzed using separate two-way ANOVAs for males and females. For males (**Figure 7E**), there was not a significant interaction between time and injury group, *F*(2.455, 54.005) = 1.704, *p* = 0.185, partial η^2^ = 0.072, ε = 0.614 (Greenhouse-Geisser); however, there was a significant main effect of injury group, *F*(1, 22) = 9.281, *p* = 0.006, partial η^2^ = 0.297, and time, *F*(2.455, 54.005) = 5.051, *p* = 0.006, partial η^2^ = 0.187, ε = 0.614 (Greenhouse-Geisser). Together, these results indicate that while all males increased the number of entries into the open areas over time, only healthy males dramatically increased the number of entries with time, making more entries than burned males. Females (**Figure 7F**) also failed to show a significant interaction between time and injury group, *F*(2.531, 55.674) = 0.310, p = 0.784, partial η^2^ = 0.014, ε = 0.633 (Greenhouse-Geisser). There was a significant main effect of time, *F*(2.531, 55.674) = 12.149, *p* < 0.001, partial η^2^ = 0.356, ε = 0.633 (Greenhouse-Geisser), indicating that there was an increase in entries into the open area over time, and this was similar for both the control and burn rats, as indicated by a non-significant main effect of injury group, *F*(1, 22) = 0.174, *p* = 0.681, partial η^2^ = 0.008. This finding suggests that, unlike males, female rats in the burn condition did not express signs of post-burn anxiety.

## Discussion

In this study, male and female rats that sustained a unilateral full-thickness paw burn with preceding traumatic stress displayed divergent, sex-specific evoked pain responses, gait dysfunction, and anxiety-like behaviors. Of the two evoked pain assessments we examined – mechanical allodynia and thermal hyperalgesia – only mechanical allodynia was present in both sexes. While both male and female rats with burns showed mechanical allodynia post-burn compared to sex-matched controls, males exhibited more severe and persistent allodynia than females. Conversely, thermal hyperalgesia was only present in males with burn injuries, relative to sex-matched controls. Females with burn injuries, however, did not demonstrate thermal hyperalgesia at any timepoint post-burn. Instead, burn-injured females showed an unexpected *resilience* to the noxious infrared probe. They demonstrated hypoalgesia, with longer latencies to lift the burned foot that lasted throughout the entire post-burn follow-up period. While there is evidence of sex differences in evoked pain responses, the expression of hypoalgesia to a noxious heat stimulus was a unique and unexpected sex difference in the sensory experience of male and female rats after the same stressors and injuries.

The mechanisms of hyperalgesia propagation and maintenance may be more dissimilar between males and females than those of allodynia propagation and maintenance, leading to sex differences in hyperalgesic behavior and treatment-induced reversal, even when allodynic behavior across sexes is similar. In a study using unilateral injections of Complete Freund’s Adjuvant (CFA) into the plantar surface of the hind paw, males and females exhibited similar severity of mechanical allodynia and thermal hyperalgesia.^28^ Treatment of this inflammatory pain with either intraperitoneal or intraplantar Δ^9^-tetrahydrocannabinol (THC) reduced allodynia similarly in both sexes, but females experienced a greater reduction in their hyperalgesia than males.^28^ This is consistent with our data, in that following the same burn injury, males and females had markedly different hyperalgesic responses, but similar allodynic responses. In previous studies using noxious experimental pain stimulation, female rats were more sensitive to cold on their hind paws and faces than males, while male rats were more sensitive to heat than females.^29^ This occurred in various experimental settings, including a nociceptive preference setup, and environments with counteracting stressors or rewards.^29^ Similarly, we found a significant difference in sensitivity to the noxious thermal stimulus between male and female rats at all timepoints, including baseline. To account for these differences, data were analyzed as a ratio of the injured (right) paw to the uninjured contralateral (left) paw for each rat.

The evoked pain behaviors tested provide a useful but limited snapshot of the rats’ pain experience, capturing only one domain of pain: reflexive, involuntary responses. The CatWalk^TM^ XT, however, captures *voluntary* movement-induced pain, which is more translational to the human experience of movement-induced pain and disability. Results found that females with burn injuries demonstrated persistent gait dysfunction relative to sex-matched controls and relative to males with burn injuries. Following burn injury, females exhibited increased guarding behavior. This was demonstrated by using a smaller area of their burned hind paw with each step, stepping more intensely on that small area, taking longer to swing the burned hind paw from step to step, and taking shorter steps with the burned hind paw. These gait changes suggest that the female rats shifted their weight heavily to the contralateral side, compared to the even left-to-right balance of the sex-matched controls. Contrarily, while males with burn injuries also demonstrated some aspects of gait dysfunction, their symptoms were less severe and did not last as long as those of females with burn injuries. In light of the reduced response to evoked pain assessments in females, the increased sensitivity to gait changes was unexpected.

Females may be more susceptible to functional limitations. After nerve growth factor (NGF) injections into the right masseter muscles of healthy male and female participants, the temporomandibular disorder (TMD)-like pain caused by the injections was more severe in females than in males.^30^ Female participants reported greater peak pain while chewing than males, and greater acute muscle soreness than males, in addition to higher muscle impairment scores and larger reductions in mechanical force tolerated on their injured masseters.^30^ These functional limitation data parallel our gait analysis data, in that movement-related activities (i.e., chewing in the TMD study, and walking/running gait analysis in our study) induced more functional limitation in females than males. These sensitive measures of gait enabled us to investigate subtle functional differences that would otherwise have remained undetectable, as these differences were not apparent to experimenters observing the rats in their home cages during routine welfare checks, or in the restrictive environment of the evoked pain behavior testing cages. Without this assessment, the females would appear healthy and otherwise unbothered by ongoing burn pain, given their lack of thermal hyperalgesia and mild mechanical allodynia relative to the males. Therefore, it is imperative that future studies employ multiple sensory testing methods to capture the range of symptom types and modalities that may be affected post-injury.

Anxiety was also assessed in this paradigm using the EZM. Here, males showed consistent signs of anxiety following the stress/burn injury model, as evidenced by diminished time spent in the open areas of the maze compared to sex-matched controls. Despite receiving the same stressors and burn injury, females did not demonstrate differences between the burn and control groups at any timepoint on the EZM. This is consistent with other pain models. In a study using unilateral sciatic nerve cuffing, both male and female C57BL/6J mice demonstrated mechanical allodynia and thermal hyperalgesia on the affected legs compared to sham mice.^31^ However, despite receiving the same nerve injury, only male cuffed mice demonstrated anxiety-and depression-like symptoms 45 days post-cuffing – cuffed males spent less time in the open areas of an EZM than cuffed females, and had longer latency to feed on a novelty suppressed feeding task.^31^ Additionally, stress may affect males and females differently. In a similar burn model, stress prior to burn injury increased sensitivity to a non-painful mechanical stimulus but not a painful thermal stimulus in females^22^ while increasing the sensitivity to both a non-painful mechanical and a painful thermal stimulus in males.^18^

While it is possible that females genuinely did not experience anxiety following the combination of stress and burn injury, it may be that the EZM is insufficient to capture female rat anxiety, in a similar fashion to the evoked pain tests being insufficient to capture the true scope of female pain and discomfort post-burn. The EZM was originally developed using male Sprague-Dawley rats,^25^ and research in large groups of healthy Wistar rats using a similar device (the Elevated Plus Maze, EPM) found a baseline difference in male versus female anxiety-like behavior.^26^ Compared to the males, females in that experiment spent more time in the open arms of the EPM, traveled a greater distance overall, and made more open arm entries than males.^26^ Their results are similar to ours using Sprague-Dawley rats in this study. As such, our results may be affected by underlying sex differences in behavior on an elevated, partially enclosed maze. However, the lack of significant differences between our male and female control groups throughout the study, as well as the lack of differences between all groups at the pre-injury baseline session, suggests that the healthy male versus female differences may be strain-specific.

This study contributes to a growing body of research examining sex differences in the experience of pain. The presence of sex differences, whether in human participants or animal models, varies depending on the type of pain stimulus used (e.g., experimentally induced) or the pain condition studied.^19^ Our work explores new techniques for capturing sex differences in pain, extending beyond traditional assessments using evoked pain responses. Further investigation is needed to elucidate the mechanisms underlying these differences. For example, gonadal hormones may contribute to the less severe mechanical allodynia and thermal hypoalgesia observed in females following burn injury, as all rats in this study were gonadally intact. In a study using a reserpine-induced model of fibromyalgia, estrogen depletion via ovariectomy (OVX) made the reserpine-induced pain worse, wherein OVX females had more severe muscle hyperalgesia than intact reserpine-treated females; conversely, estrogen supplementation via exogenous 17-β estradiol treatment reversed reserpine-induced pain in OVX females.^32^ Estradiol supplementation in intact males promotes faster recovery from chronic constriction injury (CCI)-induced mechanical allodynia compared to vehicle-treated males, and promotes recovery in intact females; without treatment, females did not meaningfully recover from allodynia over time.^33^ Taken together, these two studies suggest that estrogen may be protective when assessing evoked pain responses. Therefore, analysis of our future cohorts with estrous cycle monitoring may elucidate within-female group differences. Given these effects of estrogen on pain in the literature, there may be a subgroup of females who do become hyperalgesic post-burn, perhaps in a lower-estrogen state of the estrous cycle.

## Limitations

Despite utilizing intact female rats in our study, we did not measure or monitor the estrous cycle. As such, we cannot determine how the estrous cycle impacted our results. We did not perform any debridement or excision procedures on our burn rats, nor any grafting procedures on the burn wounds, despite these being standard procedures for human full-thickness burns. Additionally, our burn model misses some aspects of human burns present in other small animal burn models, such as hypermetabolism,^34^ given the small percent of total body surface area burned.

## Conclusions

Using a model of unilateral full-thickness paw burn with preceding traumatic stress, our study found that male and female Sprague Dawley rats displayed divergent, sex-specific evoked pain responses, gait dysfunction, and anxiety-like behavior. Future studies should make every effort to utilize multimodal assessments when testing pain across sexes. Further, future studies should examine the underlying mechanisms behind these behavioral sex differences.

## Disclosures

### Data Availability

All data are available upon request.

## Acknowledgments

The authors thank the Penn State College of Medicine’s Department of Comparative Medicine for their diligent, hard-working animal care staff and veterinarians. Without their work, complex experiments like the ones in this study would not be possible.

---

**Table S1. Pain Scale Table.** Used for post-operative monitoring of all burned rats.

## References

1. World Health Organization. Burns 2018 [accessed September 7, 2025]. Available from: https://www.who.int/news-room/fact-sheets/detail/burns.

2. CDC. (2024, September 4). 2021 *NHAMCS Results and Publications*. National Hospital Ambulatory Medical Care Survey. https://www.cdc.gov/nchs/nhamcs/publications/2021-results-publications.html

3. James SL, Lucchesi LR, Bisignano C, Castle CD, Dingels ZV, Fox JT, Hamilton EB, Henry NJ, McCracken D, Roberts NLS, Sylte DO, Ahmadi A, Ahmed MB, Alahdab F, Alipour V, Andualem Z, Antonio CAT, Arabloo J, Badiye AD, Bagherzadeh M, Banstola A, Bärnighausen TW, Barzegar A, Bayati M, Bhaumik S, Bijani A, Bukhman G, Carvalho F, Crowe CS, Dalal K, Daryani A, Nasab MD, Do HT, Do HP, Endries AY, Fernandes E, Filip I, Fischer F, Fukumoto T, Gebremedhin KBB, Gebremeskel GG, Gilani SA, Haagsma JA, Hamidi S, Hostiuc S, Househ M, Igumbor EU, Ilesanmi OS, Irvani SSN, Jayatilleke AU, Kahsay A, Kapoor N, Kasaeian A, Khader YS, Khalil IA, Khan EA, Khazaee-Pool M, Kokubo Y, Lopez AD, Madadin M, Majdan M, Maled V, Malekzadeh R, Manafi N, Manafi A, Mangalam S, Massenburg BB, Meles HG, Menezes RG, Meretoja TJ, Miazgowski B, Miller TR, Mohammadian-Hafshejani A, Mohammadpourhodki R, Morrison SD, Negoi I, Nguyen TH, Nguyen SH, Nguyen CT, Nixon MR, Olagunju AT, Olagunju TO, Padubidri JR, Polinder S, Rabiee N, Rabiee M, Radfar A, Rahimi-Movaghar V, Rawaf S, Rawaf DL, Rezapour A, Rickard J, Roro EM, Roy N, Safari-Faramani R, Salamati P, Samy AM, Satpathy M, Sawhney M, Schwebel DC, Senthilkumaran S, Sepanlou SG, Shigematsu M, Soheili A, Stokes MA, Tohidinik HR, Tran BX, Valdez PR, Wijeratne T, Yisma E, Zaidi Z, Zamani M, Zhang ZJ, Hay SI, Mokdad AH. Epidemiology of injuries from fire, heat and hot substances: global, regional and national morbidity and mortality estimates from the Global Burden of Disease 2017 study. Inj Prev. 2020 Oct;26(Supp 1):i36–i45. doi: 10.1136/injuryprev-2019-043299. Epub 2019 Dec 18. PMID: 31857422; PMCID: PMC7571358.

4. Burn Incidence & Treatment in the U.S. | ABA Burn Data. (n.d.). Retrieved January 13, 2026, from https://www.ameriburn.org/prevention/burn-incidence-and-treatment-in-the-united-states

5. Wardhan R, Fahy BG. Regional Anesthesia and Acute Pain Management for Adult Patients with Burns. J Burn Care Res. 2023 Jul 5;44(4):791–799. doi: 10.1093/jbcr/irad069. PMID: 37191659.

6. Klifto KM, Hultman CS. Pain Management in Burn Patients: Pharmacologic Management of Acute and Chronic Pain. Clin Plast Surg. 2024 Apr;51(2):267–301. doi: 10.1016/j.cps.2023.11.004. Epub 2023 Nov 23. PMID: 38429049.

7. McCann C, Watson A, Barnes D. Major burns: Part 1. Epidemiology, pathophysiology and initial management. BJA Educ. 2022 Mar;22(3):94–103. doi: 10.1016/j.bjae.2021.10.001. Epub 2021 Dec 21. PMID: 35211326; PMCID: PMC8847805.

8. Egyhazi R, Fregni F, Bravo GL, Trinh NH, Ryan CM, Schneider JC. Chronic pain following physical and emotional trauma: the station nightclub fire. Front Neurol. 2014 Jun 2;5:86. doi: 10.3389/fneur.2014.00086. PMID: 24917849; PMCID: PMC4040492.

9. Mauck MC, Smith J, Liu AY, Jones SW, Shupp JW, Villard MA, Williams F, Hwang J, Karlnoski R, Smith DJ, Cairns BA, Kessler RC, McLean SA. Chronic Pain and Itch are Common, Morbid Sequelae Among Individuals Who Receive Tissue Autograft After Major Thermal Burn Injury. Clin J Pain. 2017 Jul;33(7):627–634. doi: 10.1097/AJP.0000000000000446. PMID: 28145911.

10. Klifto KM, Dellon AL, Hultman CS. Prevalence and associated predictors for patients developing chronic neuropathic pain following burns. Burns Trauma. 2020 May 1;8:tkaa011. doi: 10.1093/burnst/tkaa011. PMID: 32377542; PMCID: PMC7192663.

11. Mauck MC, Shupp JW, Williams F, Villard MA, Jones SW, Hwang J, Smith J, Karlnoski R, Smith DJ, Cairns BA, McLean SA. Hypertrophic Scar Severity at Autograft Sites Is Associated With Increased Pain and Itch After Major Thermal Burn Injury. J Burn Care Res. 2018 Jun 13;39(4):536–544. doi: 10.1093/jbcr/irx012. PMID: 29596686; PMCID: PMC7189975.

12. Van Loey NEE, de Jong AEE, Hofland HWC, van Laarhoven AIM. Role of burn severity and posttraumatic stress symptoms in the co-occurrence of itch and neuropathic pain after burns: A longitudinal study. Front Med (Lausanne). 2022 Oct 12;9:997183. doi: 10.3389/fmed.2022.997183. PMID: 36314001; PMCID: PMC9596796.

13. Malenfant A, Forget R, Papillon J, Amsel R, Frigon JY, Choinière M. Prevalence and characteristics of chronic sensory problems in burn patients. Pain. 1996 Oct;67(2-3):493–500. doi: 10.1016/0304-3959(96)03154-5. PMID: 8951946.

14. Panayi AC, Heyland DK, Stoppe C, Jeschke MG, Didzun O, Matar D, Tapking C, Palackic A, Bliesener B, Harhaus L, Knoedler S, Haug V, Bigdeli AK, Kneser U, Orgill DP, Hundeshagen G. The long-term intercorrelation between post-burn pain, anxiety, and depression: a post hoc analysis of the “RE-ENERGIZE” double-blind, randomized, multicenter placebo-controlled trial. Crit Care. 2024 Mar 22;28(1):95. doi: 10.1186/s13054-024-04873-8. PMID: 38519972; PMCID: PMC10958907.

15. Klifto KM, Yesantharao PS, Lifchez SD, Dellon AL, Hultman CS. Chronic Nerve Pain after Burn Injury: An Anatomical Approach and the Development and Validation of a Model to Predict a Patient’s Risk. Plast Reconstr Surg. 2021 Oct 1;148(4):548e–557e. doi: 10.1097/PRS.0000000000008315. PMID: 34550938.

16. Summer GJ, Puntillo KA, Miaskowski C, Green PG, Levine JD. Burn injury pain: the continuing challenge. J Pain. 2007 Jul;8(7):533–48. doi: 10.1016/j.jpain.2007.02.426. Epub 2007 Apr 16. PMID: 17434800.

17. Lewis J, Pride LC, Lee S, Anwaegbu O, Tabukumm NN, Patel MM, Lee WC. Examining the role of post-traumatic stress disorder, chronic pain and opioid use in burn patients: A multi-cohort analysis. Scars Burn Heal. 2024 Dec 11;10:20595131241288298. doi: 10.1177/20595131241288298. Erratum in: Scars Burn Heal. 2025 Sep 24;11:20595131251376836. doi: 10.1177/20595131251376836. PMID: 39665049; PMCID: PMC11632867.

18. Nyland JE, McLean SA, Averitt DL. Prior stress exposure increases pain behaviors in a rat model of full thickness thermal injury. Burns. 2015 Dec;41(8):1796–1804. doi: 10.1016/j.burns.2015.09.007. Epub 2015 Oct 1. PMID: 26432505.

19. Fillingim RB, King CD, Ribeiro-Dasilva MC, Rahim-Williams B, Riley JL 3rd. Sex, gender, and pain: a review of recent clinical and experimental findings. J Pain. 2009 May;10(5):447–85. doi: 10.1016/j.jpain.2008.12.001. PMID: 19411059; PMCID: PMC2677686.

20. Rosen SF, Ham B, Haichin M, Walters IC, Tohyama S, Sotocinal SG, Mogil JS. Increased pain sensitivity and decreased opioid analgesia in T-cell-deficient mice and implications for sex differences. Pain. 2019 Feb;160(2):358–366. doi: 10.1097/j.pain.0000000000001420. PMID: 30335680.

21. McGrath, K., Mauck, C., M., Linnstaedt, S., Sefton, C., Shan, Y., Cairns, B., & McLean, S. (2019, October 19). Mechanistic Insights into Sex Differences in Chronic Pain Following Major Thermal Burn Injury. F1046 / 7 - Mechanistic Insights into Sex Differences in Chronic Pain Following Major Thermal Burn Injury. Anesthesiology 2019, Session F02 -- Featured Abstracts II. https://www.abstractsonline.com/pp8/#!/6832/presentation/6831

22. Strain MM, Tongkhuya S, Wienandt N, Alsadoon F, Chavez R, Daniels J, Garza T, Trevino AV, Wells K, Stark T, Clifford J, Sosanya NM. Exploring combat stress exposure effects on burn pain in a female rodent model. BMC Neurosci. 2022 Dec 6;23(1):73. doi: 10.1186/s12868-022-00759-z. PMID: 36474149; PMCID: PMC9724288.

23. Sosanya NM, Garza TH, Stacey W, Crimmins SL, Christy RJ, Cheppudira BP. Involvement of brain-derived neurotrophic factor (BDNF) in chronic intermittent stress-induced enhanced mechanical allodynia in a rat model of burn pain. BMC Neurosci. 2019 Apr 24;20(1):17. doi: 10.1186/s12868-019-0500-1. PMID: 31014242; PMCID: PMC6480655.

24. Garrick JM, Costa LG, Cole TB, Marsillach J. Evaluating Gait and Locomotion in Rodents with the CatWalk. Curr Protoc. 2021 Aug;1(8):e220. doi: 10.1002/cpz1.220. PMID: 34370398; PMCID: PMC8363132.

25. Shepherd JK, Grewal SS, Fletcher A, Bill DJ, Dourish CT. Behavioural and pharmacological characterisation of the elevated “zero-maze” as an animal model of anxiety. Psychopharmacology (Berl). 1994 Sep;116(1):56–64. doi: 10.1007/BF02244871. PMID: 7862931.

26. Knight P, Chellian R, Wilson R, Behnood-Rod A, Panunzio S, Bruijnzeel AW. Sex differences in the elevated plus-maze test and large open field test in adult Wistar rats. Pharmacol Biochem Behav. 2021 May;204:173168. doi: 10.1016/j.pbb.2021.173168. Epub 2021 Mar 5. PMID: 33684454; PMCID: PMC8130853.

27. Sorge RE, Martin LJ, Isbester KA, Sotocinal SG, Rosen S, Tuttle AH, Wieskopf JS, Acland EL, Dokova A, Kadoura B, Leger P, Mapplebeck JC, McPhail M, Delaney A, Wigerblad G, Schumann AP, Quinn T, Frasnelli J, Svensson CI, Sternberg WF, Mogil JS. Olfactory exposure to males, including men, causes stress and related analgesia in rodents. Nat Methods. 2014 Jun;11(6):629–32. doi: 10.1038/nmeth.2935. Epub 2014 Apr

28. PMID: 24776635.

28. Craft RM, Kandasamy R, Davis SM. Sex differences in anti-allodynic, anti-hyperalgesic and anti-edema effects of Δ(9)-tetrahydrocannabinol in the rat. Pain. 2013 Sep;154(9):1709–1717. doi: 10.1016/j.pain.2013.05.017. Epub 2013 May 15. PMID: 23707295.

29. Vierck CJ, Acosta-Rua AJ, Rossi HL, Neubert JK. Sex differences in thermal pain sensitivity and sympathetic reactivity for two strains of rat. J Pain. 2008 Aug;9(8):739–49. doi: 10.1016/j.jpain.2008.03.008. Epub 2008 May 16. PMID: 18486556; PMCID: PMC2547088.

30. Schabrun SM, Si E, Millard SK, Chiang AKI, Chen S, Chowdhury NS, Seminowicz DA. Intramuscular injection of nerve growth factor as a model of temporomandibular disorder: nature, time-course, and sex differences characterising the pain experience. Neurobiol Pain. 2023 Jan 13;13:100117. doi: 10.1016/j.ynpai.2023.100117. PMID: 36687467; PMCID: PMC9852786.

31. Cardenas A, Caniglia J, Keljalic D, Dimitrov E. Sex differences in the development of anxiodepressive-like behavior of mice subjected to sciatic nerve cuffing. Pain. 2020 Aug;161(8):1861–1871. doi: 10.1097/j.pain.0000000000001875. Epub 2020 Mar 20. PMID: 32701845; PMCID: PMC7502469.

32. Hernandez-Leon A, De la Luz-Cuellar YE, Granados-Soto V, González-Trujano ME, Fernández-Guasti A. Sex differences and estradiol involvement in hyperalgesia and allodynia in an experimental model of fibromyalgia. Horm Behav. 2018 Jan;97:39–46. doi: 10.1016/j.yhbeh.2017.10.011. Epub 2017 Nov 8. PMID: 29080671.

33. Vacca V, Marinelli S, Pieroni L, Urbani A, Luvisetto S, Pavone F. 17beta-estradiol counteracts neuropathic pain: a behavioural, immunohistochemical, and proteomic investigation on sex-related differences in mice. Sci Rep. 2016 Jan 8;6:18980. doi: 10.1038/srep18980. PMID: 26742647; PMCID: PMC4705539.

34. Vinaik R, Aijaz A, Jeschke MG. Small animal models of thermal injury. Methods Cell Biol. 2022;168:161–189. doi: 10.1016/bs.mcb.2021.12.014. Epub 2022 Jan 10. PMID: 35366981; PMCID: PMC10107476.

