## Supplemental Table 1 for "Pronounced Sex Differences in Evoked and Spontaneous Pain Assessments Following Full-Thickness Traumatic Burn Injury in Male and Female Sprague Dawley Rats"

| **Assessment Score =** | **-0-** | **-1-** | **-2-** | **-3-** |
|---|---|---|---|---|
| **Attitude and Posture (all pain models)** | *Alert and not hunched* | *Not Alert or hunched* | *Not alert and hunched* | *Not responsive to stimuli* |
| **Gait and Movement (all pain models)** | *Active* | *Somewhat inactive* | *Completely inactive* | *Lying on side* |
| **Appetite (all pain models)** | *Eating and drinking normally No weight loss* | *Reduced eating or drinking Or weight loss of less than 10% compared to control* | *Not eating or drinking Or weight loss between 10-20% compared to control* | *Not eating or drinking for >3 days Or weight loss greater than 20% compared to control* |
| **Elimination (all pain models)** | *Normal* | *Softer than normal* | *Diarrhea* | *Diarrhea > 3 days* |

**Table S1. Pain Scale Table.** Used for post-operative monitoring of all burned rats.
